# Maternal H3.3-Mediated Paternal Genome Reprogramming Contributes to Minor Zygotic Genome Activation

**DOI:** 10.1101/2023.11.07.566007

**Authors:** Jiaming Zhang, Xuanwen Li, Hongdi Cui, Songling Xiao, Entong Song, Ming Zong, Shukuan Ling, Zev Rosenwaks, Shaorong Gao, Xiaoyu Liu, Qingran Kong, Duancheng Wen

**Author notes:** Correspondence: Qingran Kong, Xiaoyu Liu, Duancheng Wen. These authors contributed equally to the work (Jiaming Zhang, Xuanwen Li).

## Abstract

In mice, zygotic genome activation (ZGA) is initiated at the late-one-cell stage, accompanied by an extensive incorporation of the histone variant H3.3 into the parental genomes. However, it is unclear how H3.3 engages in the onset of ZGA. Here, using the H3.3B-HA-tagged mouse model, we found that the paternal and maternal genomes are activated asynchronously. Paternally expressed H3.3 begins deposition on the zygotic genome at the early two-cell stage, whereas the deposition of maternally expressed H3.3 is delayed until the four-cell stage. Oocyte-stored maternal H3.3 (mH3.3) is crucial for cleavage development and minor ZGA. Deposition of mH3.3 on the paternal genome occurs globally during the protamine-to-histone transition but shows preferential enrichment at CpG-rich TSSs after the initial round of DNA replication. Depletion of mH3.3 can lead to a loss of H3K27ac, resulting in minor ZGA failure and early embryonic arrest. Mechanistically, mH3.3 deposition on the sperm genome removes repressive histone modifications, promotes the establishment of active histone modifications, and in turn enables the initiation of minor ZGA from the paternal genome. Our study highlights the pivotal role of mH3.3 in paternal genome reprogramming and minor ZGA initiation.

## Introduction

Embryogenesis begins with a single cell, which is formed through the fusion of the egg’s cytoplasm and the haploid genomes from both parents, giving rise to a unified diploid zygotic genome (1). In the beginning, the development is mainly governed by maternal factors present in the egg’s cytoplasm, while the zygotic genome remains inactive. As development progresses, both the maternal and paternal genomes undergo extensive epigenetic reprogramming, facilitating zygotic genome activation (ZGA) (2). In mice, ZGA is orchestrated through two waves of de novo gene expression that activate a specific set of genes. The initial minor wave occurs at the S phase of the one-cell stage zygote and the G1 stage of the early 2-cell embryo. The major wave, on the other hand, happens at the mid to late two-cell stage, following the second round of DNA replication (2–4). The minor ZGA must precede the major ZGA for the successful execution of the maternal-to-zygotic transition. The timely onset of minor ZGA is critical for preimplantation development to progress beyond the two-cell stage (5).

Various facets of epigenetic regulation, such as DNA methylation, histone modifications, and chromatin remodeling, have been associated with minor ZGA. The histones around which DNA is wrapped undergo various post-translational modifications such as methylation, acetylation, and phosphorylation. These modifications can influence chromatin structure and gene expression. For instance, histone H3 lysine 4 trimethylation (H3K4me3) and histone H3 lysine 27 trimethylation (H3K27me3) are known to be involved in gene activation and repression, respectively, and might play a role in ZGA (6–10). Epigenetic marks inherited from the sperm and egg might also play a role in regulating minor ZGA. These epigenetic marks can affect gene expression and might be important for the differential expression of maternal and paternal alleles during minor ZGA (2, 11–13). Despite these insights, several critical questions remain unanswered: 1) What are the specific epigenetic signatures in the parental genomes that allow minor ZGA genes to be activated? 2) How are these minor ZGA genes activated after fertilization?

Once the sperm enters the egg during fertilization, the paternal DNA must be reorganized to facilitate the subsequent steps of embryonic development. This reorganization involves the replacement of protamines with histones. The protamines are removed from the sperm chromatin, and the DNA is repackaged with maternal histones supplied by the oocyte. The protamine-to-histone transition is essential for the proper reprogramming of the paternal genome (14–16). It ensures that the chromatin of the paternal DNA adopts a configuration that is compatible with the transcriptional and regulatory requirements of the early embryo. This transition also plays a role in epigenetic reprogramming, as new histones can carry various modifications that influence gene expression. However, the exact mechanisms of this process and roles of these modification patterns in the regulation of ZGA remain to be fully elucidated.

The H3 histone variant, H3.3, is encoded by two genes, *h3f3a* (H3.3A) and *h3f3b* (H3.3B), both of which generate an identical protein product (17, 18). H3.3A and H3.3B are expressed redundantly during oogenesis and continue through the preimplantation stages and are capable of compensating for each other during embryonic development (19). This functional redundancy is further supported by the observation that the deletion of either gene does not result in any noticeable developmental phenotype in mice (20, 21). H3.3 is crucial for the reprogramming of paternal and maternal genomes during fertilization (19, 22–24), owing to its unique characteristic of being incorporated into chromatin independent of DNA replication. In contrast, its canonical counterparts H3.1 and H3.2 are deposited into chromatin in a DNA replication-dependent manner (25). This feature distinguishes H3.3 as the sole H3 variant that forms nucleosomes and facilitates genome remodeling after fertilization and prior to the first round of DNA replication.

The genomes of both sperm and oocyte are enriched with H3.3 protein, referred to as sH3.3 (sperm genome-associated H3.3) and oH3.3 (oocyte genome-associated H3.3), respectively. Furthermore, H3.3 mRNAs, termed mH3.3, are present in the oocyte cytoplasm and enable the synthesis of H3.3 protein immediately after fertilization. The zygotic genome starts transcribing H3.3 at the early 2-cell stage, and this zygotically expressed H3.3 is denoted as zH3.3 (26). We previously reported that sH3.3 is rapidly removed from the sperm genome post fertilization (23), while mH3.3 is integrated into the paternal genome as early as 1 hour post-fertilization, with detectable levels persisting until the morula stage (23). Our studies (19, 23), along with that of others (21, 22, 24), have underscored the critical role of mH3.3 in fertilization and early embryonic development. A major challenge arises due to the coexistence of distinct sources of H3.3 in early embryos, namely oH3.3, mH3.3, and zH3.3. This concurrent presence complicates our ability to precisely track the individual dynamics of these H3.3 variants, which may have distinct roles in embryonic development. Consequently, several questions remain unanswered, particularly regarding the specific deposition of mH3.3 into the paternal genome, and how this deposition contributes to the reprogramming of the paternal genome and the initiation of ZGA.

In this study, we investigated the dynamic deposition of mH3.3 on the paternal genome and its potential function during fertilization, employing micromanipulation and low-input ChIP-seq in H3.3B-HA reporter mice. We found that mH3.3 plays a pivotal role in paternal genome reprogramming and the onset of minor ZGA.

## Results

### Dynamic Profiling of H3.3 Deposition in Mouse Preimplantation Embryos

To track endogenous H3.3 gene expression, we developed a reporter system by knocking-in a small C-terminal HA tag into the last coding exon of the H3.3B gene. This HA tag was accompanied by an internal ribosomal entry site (IRES) and a distinct translation sequence for either EYFP or mCherry, serving as gene expression indicators (26). ChIP-seq analyses performed on cells from H3.3B-HA tagged mice using an HA antibody unveiled genomic enrichment patterns echoing those detected with H3.3 antibodies (25, 27), underlining the robustness of this reporter system in tracing H3.3 deposition.

To investigate the dynamics of H3.3 deposition in early mouse embryos, we conducted ultra-low-input native chromatin immunoprecipitation sequencing (ULI-NChIP-seq) (28) using preimplantation embryos harvested from H3.3B-HA reporter mice. Embryos expressing H3.3B-HA maternally (WT♂ × H3.3B-HA ♀) and those expressing the paternal allele of H3.3B-HA (H3.3B-HA ♂ × WT ♀) were collected separately (**Fig. 1A**). It is important to note that H3.3B-HA in the embryos from the maternal series (referred to as maH3.3) encompasses oH3.3B-HA, mH3.3B-HA, and zH3.3B-HA. In contrast, H3.3B-HA in the embryos from the paternal series (denoted as paH3.3) exclusively contains zH3.3B-HA, which is expressed from the paternal allele.

**Figure 1.**
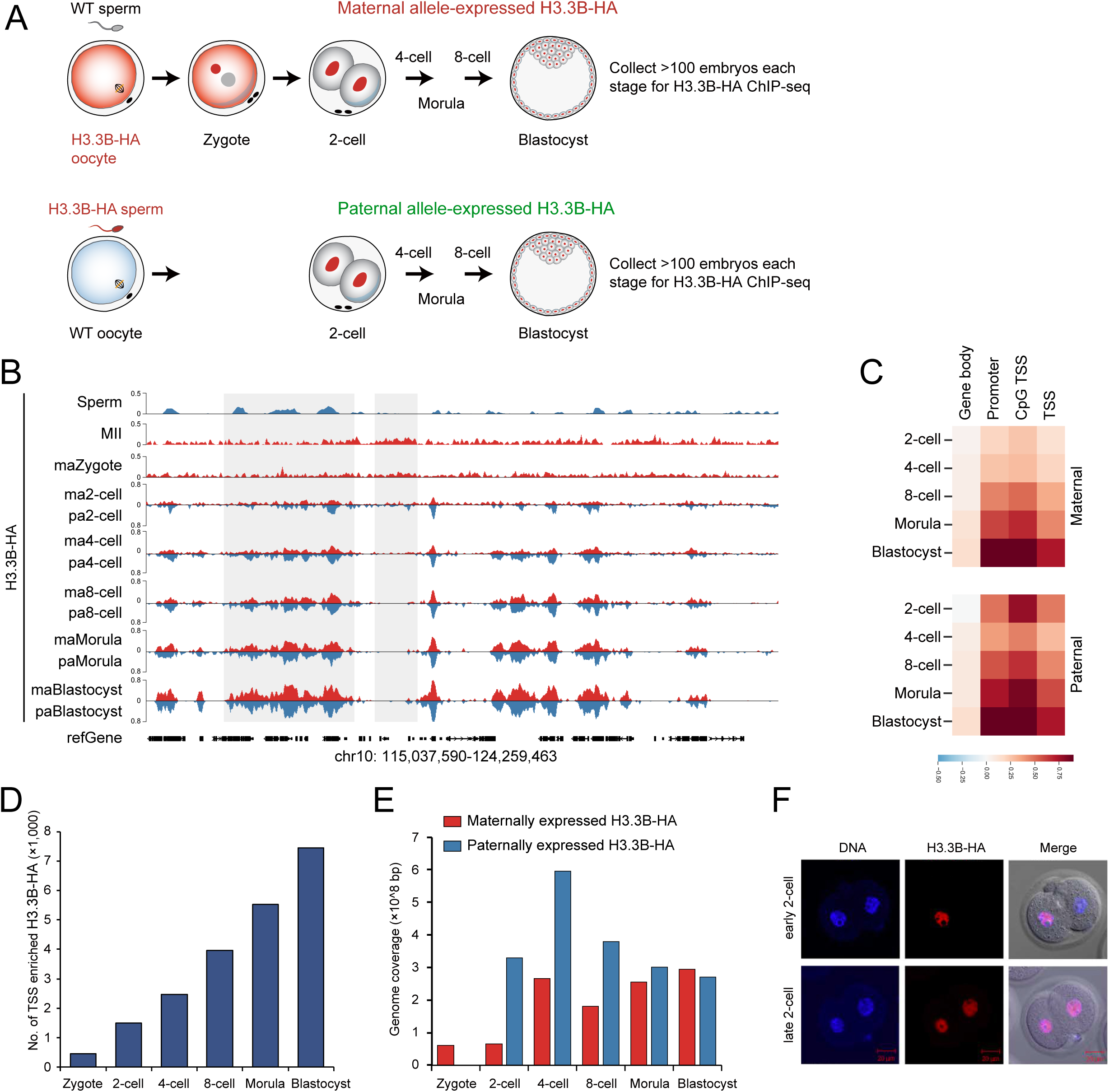
Dynamic Profiling of H3.3 Deposition in Mouse Preimplantation Embryos. (A) Schematic illustrations of the experiments. Preimplantation embryos at different stages were collected (>100 embryos each stage) for H3.3B-HA ChIP-seq. Two sets of embryos were collected to measure deposition differences between maternally and paternally expressed H3.3 in early embryonic development. (B) The genome browser view showing enrichment of maternal and paternal H3.3 signals in mouse gametes and early embryos. Signals represent the log2-transformed H3.3/input ratio. The maternal and paternal H3.3 enrichments are indicated in red and blue, respectively. m, maternal H3.3 enrichment in the indicated embryos. p, paternal H3.3 enrichment in the indicated embryos. (C) Heatmap of H3.3 enrichment from maternal and paternal series for various annotated elements. (D) Number of TSSs with >2-fold increase in H3.3 deposition compared with that in oocytes at different developmental stages. (E) Total genome coverage for maternal and paternal samples. (F) Immunostaining of 2-cell stage embryos (H3.3B-HA sperm and WT oocyte) revealed strong H3.3B-HA signals originating from the paternal allele at the early and late 2-cell stage. This suggests that H3.3 from the paternal allele can be activated at the early 2-cell stage.

The H3.3B-HA ULI-NChIP data from mouse embryos exhibited remarkable reproducibility (**Fig. S1A**). Principal component analysis (PCA) revealed that while maH3.3 and paH3.3 enrichment patterns largely aligned during pre-implantation development, a notable difference was evident at the 2-cell stage. Intriguingly, maH3.3 enrichment traced a consistent developmental curve for mouse preimplantation embryos, a trend that was disrupted in paH3.3 specifically at the 2-cell stage (**Fig. S1B**). This highlights the critical influence of the maternal H3.3 effect, encompassing mH3.3 and oH3.3, in reprogramming the parental genome of mouse pre-implantation embryos.

While H3.3 enrichment patterns in oocytes and sperm exhibit significant differences, the canonical H3.3B-HA peaks from both the maternal and paternal series show a notable resemblance from the 2-cell stage onward (**Fig. 1B and S1C**). These typical H3.3B-HA enrichment patterns are predominantly observed in regulatory regions and CpG-rich transcription start sites (TSSs) (**Fig. 1B, 1C**). Interestingly, while these peaks align with H3.3 peaks in the sperm genome, they don’t match those in the oocyte genome (**Fig. 1B**). This suggests that the retention of H3.3 in the sperm genome might be linked to genes crucial for early development. The number of TSSs with H3.3B-HA enrichment progressively increases from the zygote stage through to the blastocyst stage, suggesting an escalating reprogramming of genes in tandem with developmental progression during the early embryonic stages (**Figure 1D**).

An analysis of the total base pairs covered by H3.3B-HA peaks showed that the H3.3B-HA coverage in the maternal series remained consistent between zygotes and 2-cell embryos (**Fig. 1E**). However, a pronounced four-fold increase in coverage was observed by the 4-cell stage, with this heightened level persisting from the 8-cell stage through to the blastocyst stages (**Fig. 1E**). The consistently low H3.3B-HA coverage in zygotes and 2-cell embryos, followed by the marked rise at the 4-cell stage, suggests the initiation of H3.3B-HA expression and deposition from the maternal genome begins at the 4-cell stage. Conversely, the paternal series displayed a prominent increase in H3.3B-HA deposition at the 2-cell stage when compared to the maternal series (**Fig. 1E**). This is further supported by the immunostaining of paH3.3B-HA and the asynchronous expression seen exclusively in the early 2-cell stage embryos (**Fig. 1F**). Collectively, our data suggest that H3.3B activation in the paternal genome commences at the early 2-cell stage, whereas its initiation in the maternal genome is deferred until the 4-cell stage. This asynchronous activation of the H3.3B gene between paternal and maternal genomes underscores a differential reprogramming of the parental genomes, with the paternal genome becoming active earlier than the maternal genome during ZGA.

### Deposition of mH3.3 on the Paternal Genome Post-Fertilization

In a prior study, we found that mH3.3 plays a crucial role in activating the paternal genome (23). Building on this discovery, we sought to examine the precise deposition of mH3.3 on the paternal genome after fertilization. To achieve this, we enucleated oocytes from H3.3B-HA reporter mice and injected two wildtype (WT) sperm heads to generate diploid androgenetic embryos. At specified intervals, these embryos were subjected to ChIP-seq analysis with an HA antibody, aiming to pinpoint the dynamic mH3.3 deposition on the paternal genome (**Fig. 2A**). A major event in the reprogramming of the paternal genome following fertilization is the protamine-to-histone transition (14–16, 29). During this transition, mH3.3 is the sole histone H3 variant capable of replacing protamines in the sperm genome, facilitating the assembly of nucleosomes before the first round of DNA replication (**Fig. 2A**).

**Figure 2.**
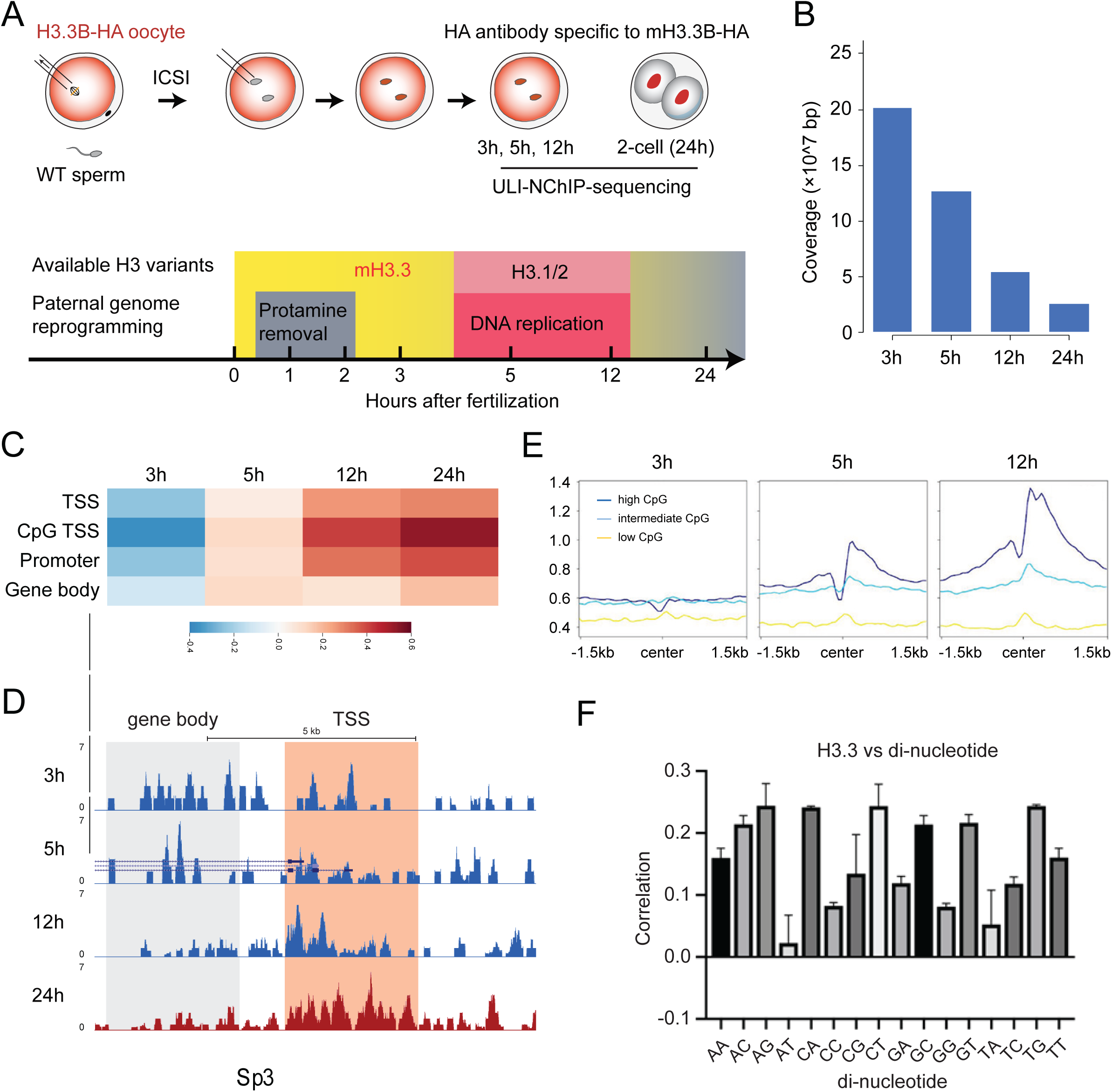
Tracking mH3.3 deposition on the paternal genome post fertilization. (A) (A) Schematic illustration of the generation of androgenetic embryos. Embryos at different time points were collected to track mH3.3 deposition on the paternal genome using HA antibody. Major events in paternal genome reprogramming were shown below. (B) Coverage of the paternal genome by mH3.3 based on H3.3B-HA ULI-NChIP data. (C) mH3.3 enrichment at different annotations in the paternal genome after fertilization. (D) Tracks of mH3.3 enrichment on the Sp3 gene (a transcription factor that binds to the GC- and GT-box regulatory elements in target genes). (E) Deposition of H3.3 around the TSS was strongly dependent on the CpG content of the TSS region. Dynamic changes of mH3.3 patterns in androgenetic embryos. (F) Correlation of dinucleotides with H3.3 replicated broad peaks.

We examined the genome-wide coverage by mH3.3B-HA and observed that the coverage peaked 3 h post-fertilization, then steadily diminished from 3 to 24 h (during the 2-cell stage) in androgenetic embryos (**Fig. 2B**). We employed fold change over input as a metric for evaluating H3.3B-HA enrichment. We found that approximately 75% of these windows contained at least one locus with a fold change greater than 1, and around 50% of the windows had at least one locus with a fold change exceeding 2 (**Fig. S2B**). Therefore, our data suggests that mH3.3 deposition is dispersed throughout the entire paternal genome before 12 h timepoints.

To pinpoint the location of mH3.3 deposition on the paternal genome, we analyzed the percentage of peak coverage across five primary functional genome annotations – intergenic regions, introns, coding sequences (CDS), untranslated regions (UTR), and transcription start sites (TSS). We observed that the majority of peaks were situated in intergenic regions (46%) and introns (42%) at the 3 h timepoint (**Figs. 2C and S2C**). Although the percentage of peaks in introns remained consistent over time, there was a decline from 46% to 25% in intergenic regions at 24 h timepoint (**Fig. S2C**). Conversely, the coverage of mH3.3 at the TSS saw a fivefold increase, rising from 2% at 3 h to 10% at 24 h timepoints (**Fig. S2C**). This implies that the TSS and intergenic regions are the most dynamic sites for mH3.3B-HA deposition during fertilization. Additionally, 7,163, 9,000, and 8,768 genes displayed enrichment in mH3.3 at the TSS (with fold change greater than 2) at 3, 5, and 12 h respectively. However, only 1,120 genes exhibited overlap at all these timepoints (**Fig. S2D**). This suggests that the deposition of mH3.3B-HA at the TSS is highly dynamic and potentially stochastic following fertilization and before the first embryonic division.

Since DNA replication commences as early as 4 h post-fertilization on the paternal genome (30), canonical H3 variants H3.1/H3.2 have the potential to be loaded during this period to form nucleosomes, thereby competing with mH3.3 deposition (**Fig. 2A**). Consistent with this, we observed that the genome coverage by mH3.3B-HA on the paternal genome progressively decreases from 3 to 24 h (**Fig. 2B**), suggesting a possible dilution of mH3.3B-HA due to the incorporation of canonical H3.1/3.2 histones after DNA replication. In evaluating the enrichment across different types of functional genome annotations, we found that the deposition of mH3.3B-HA on the paternal genome was widespread among the annotations at 3 and 5 h (**Fig. 2C, S2B-S2C**). However, after DNA replication (at 12 and 24 h), it predominantly occurred at promoters, specifically at CpG-rich TSSs and CpG islands (**Figs. 2C-2E**). We performed k-means clustering and found that the mH3.3B-HA peaks can be clustered into two major groups (**Fig. S2E**). The cluster I peaks were mainly sperm-specific and lost after fertilization. And the widespread mH3.3B-HA signals of the cluster II were established following fertilization. Most of the cluster II covered genes were related to the regulation of transcription and chromosome organization (**Fig. S2F**). Collectively, our observations suggest that the deposition of mH3.3 on the paternal genome initiates in a widespread manner across the genome, but following the first round of DNA replication, it becomes preferentially enriched at CpG-rich TSSs, which may contribute to the activation of paternal genes.

To understand this preferential CpG enrichment, we performed single and dinucleotide frequency analyses to determine the genome-wide nucleotide preferences for H3.3B-HA enrichment. There was an increase in the frequency of G/C nucleotides at 12, 24, and 48 h and a decrease in the frequency of A/T nucleotides at 12 and 48 h (4-cell stage) (**Fig. S2G**). In dinucleotide analysis, we found a depletion of AT or TA with H3.3B-HA enrichment (**Fig. 2F**). Interestingly, CG dinucleotide frequency significantly increased at 5 and 12 h, and a relatively stable frequency of GC dinucleotides was observed at 3, 5, 12, and 48 h (**Fig. S2H**). Therefore, our observations suggest that AT, CG, and TA contribute to H3.3 positioning before the first round of DNA replication and that CG contributes to H3.3 enrichment throughout the post-replication period.

### Depletion of mH3.3 Leads to Two-cell Arrest and Failure of Minor ZGA

Our prior research indicates that mH3.3 is crucial for both the activation of the paternal genome and early embryonic development (19, 23). To investigate the functional mechanism of mH3.3 in regulating early embryonic development, we depleted mH3.3 by injecting siRNA and morpholino targeting both *h3f3a* and *h3f3b* into MII oocytes, followed by intracytoplasmic sperm injection (ICSI) (**Fig. 3A**). The successful knockdown of mH3.3 was confirmed by immunostaining at the zygote stage using an H3.3-specific antibody (**Fig. 3B**). The H3.3 signal exhibited significant reduction in the injected zygotes, as evidenced by the CUT&Tag method after spike-in normalization (**Fig. 3C**). Notably, the development of mH3.3-depleted embryos was severely impaired, with most embryos arresting at the two-cell stage (**Fig. 3D**). These observations underscore the pivotal role of mH3.3 in early pre-implantation embryonic development.

**Figure 3.**
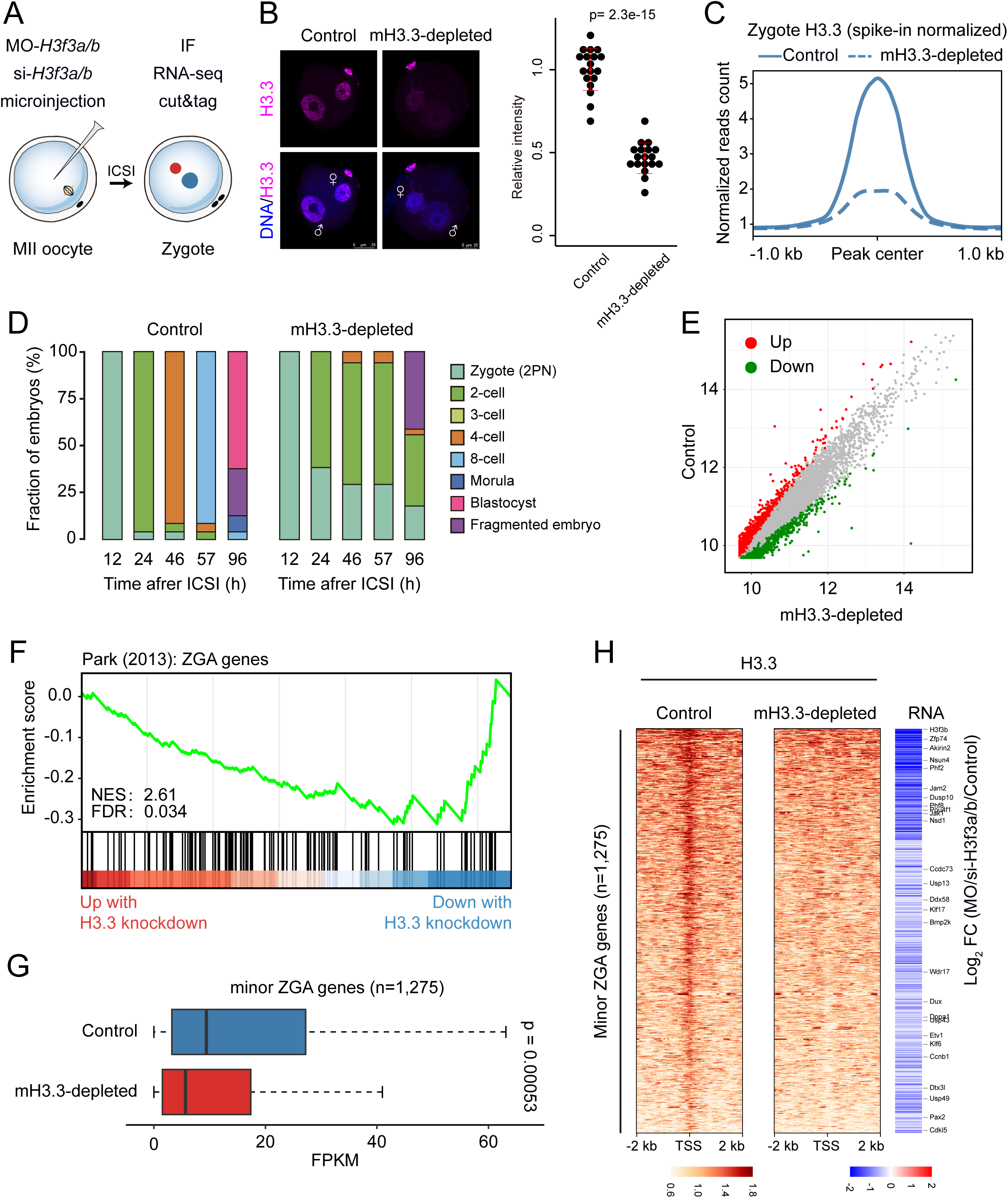
Deposition of mH3.3 is required minor ZGA gene activation. (A) Schematic presentation of the mH3.3 knockdown experimental protocol. Results of immunofluorescence (B) and CUT&Tag analysis (C) of mH3.3-depleted zygotes, validating the mH3.3 knockdown efficiency using H3.3-specific antibody. (D) Stacked bar plots showing fraction of mouse embryos at the different developmental stages by mH3.3 knockdown. (E) Scatter plots showing the whole transcriptome changes in control and mH3.3-depleted zygotes. (F) Gene set enrichment analysis (GSEA) of ZGA genes showing preferential downregulation in mH3.3-depleted embryos. (G) boxes plot showing the minor ZGA genes (n=1,275) were significantly downregulated in mH3.3-depleted embryos. (H) Heatmap (left) showing H3.3 signals ranked by their relative changes by mH3.3 knockdown. Heatmap (right) showing the downregulation of minor ZGA genes in mH3.3-depleted embryos. Mean values of two biological replicates were scaled and are represented as Z scores.

Given that most mH3.3-depleted embryos were arrested at the 2-cell stage (**Fig. 3D**), we hypothesized that the knockdown of mH3.3 might impact ZGA gene activation. To explore this, we performed RNA-seq analysis on both control and mH3.3-depleted zygotes. The transcriptome analysis identified that over 2,000 genes were downregulated in mH3.3-depleted embryos (**Fig. 3E-F**). Notably, a significant portion of the downregulated genes belonged to the category of previously reported ZGA genes (2), with a large subset being minor ZGA genes (1,275 genes, designated mH3.3-depedent minor ZGA genes) including the *h3f3b* (H3.3B) and *Dux* (**Fig. 3F-H**). These downregulated genes also exhibited a marked decrease in H3.3 deposition at their TSSs (**Fig. 3C, 3H**), suggesting the activation of these minor ZGA genes depends on mH3.3 deposition during fertilization. Collectively, our findings strongly suggest that mH3.3 deposition is associated with the activation of minor ZGA genes post-fertilization.

### Establishment of mH3.3K27ac in the Paternal Genome is Essential for Minor ZGA

It is well-documented that the transition of genes from a silenced state to an active state necessitates substantial epigenetic reprogramming within the promoter regions (31–34). In this context, modifications to histone H3 are considered to play significant roles, as postulated by the “histone code” hypothesis (35–37). Following this understanding, we proceeded to examine which histone modifications are influenced by the deposition of mH3.3 on these minor ZGA genes. Accordingly, we conducted an analysis to evaluate the correlations between mH3.3 and H3 modifications at the zygotic stage. Remarkably, the deposition pattern of mH3.3 mirrored that of H3K27ac, exhibiting similar distribution profiles and displaying a robust correlation with this modification (r=0.78) in zygotes (**Figs. 4A; and S3A**). Indeed, H3K27ac was prominently enriched at genomic elements that coincided with regions enriched for mH3.3 (**Fig. 4B**). In line with this, recent studies have emphasized the vital role of H3K27ac in regulating ZGA (38–41). Given this information, we postulated a functional connection between mH3.3, H3K27ac, and minor ZGA. To investigate this, we first assessed H3K27ac enrichment in mH3.3-depleted embryos using immunofluorescence staining and observed a significant decrease in enrichment (**Fig. 4C**). Furthermore, we also observed substantial reductions in H3K27ac signals at the TSS regions in mH3.3-depleted embryos (**Figs. 4D**), suggesting that the deposition of mH3.3 is intrinsically associated with the establishment of H3K27ac at promoter regions in zygotes post-fertilization.

**Figure 4.**
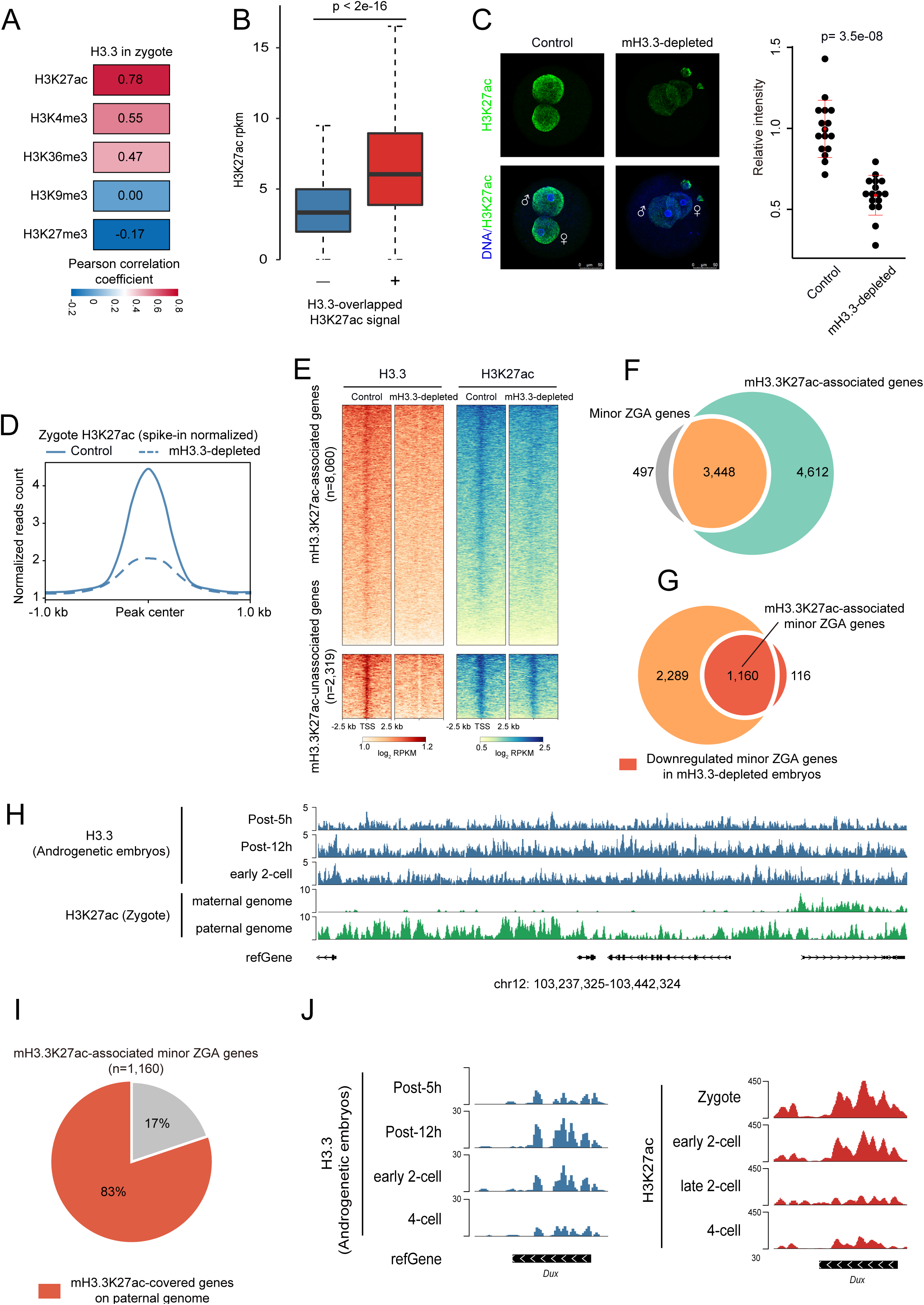
Establishment of mH3.3K27ac at promoters is essential for minor ZGA. (A) Global spearman correlation of H3.3 with other epigenetic markers at zygote stage. Bin size: 1 kb. (B) Box plots showing the comparison of H3K27ac rpkm at H3.3 and non-H3.3 enriched regions. (C) Representative confocal images of control and mH3.3-depleted embryos stained with H3K27ac antibody (left). Quantification of fluorescence intensity in control and mH3.3-depleted embryos. Each dot represents a single nucleus (right). (D) Metaplot of H3K27ac signals (spike-in normalized, Z-score normalized) in control and mH3.3-depleted zygotes. (E) Heatmap analysis of H3.3 and H3K27ac signals at promoters (n=10,379) ranked by the change in H3K27ac enrichment after deletion of mH3.3. (F) Venn diagrams showing the overlap of minor ZGA genes and H3.3K27ac-associated genes defined in Fig. 3F. (G) Venn diagrams showing the overlap of the intersection genes in Fig. 3G and downregulated minor ZGA genes in mH3.3-depleted embryos. (H) The genome browser view shows the paternal enrichments of H3K27ac exhibited high correlation with those of mH3.3 on the paternal genome. (I) Pie chart showing the fraction of mH3.3K27ac-associated minor ZGA genes covered by mH3.3K27ac on paternal genome. (J) Genome browser view of H3.3 and H3K27ac signals at the *Dux* locus.

Next, we investigated how mH3.3K27ac regulates gene expression during ZGA. We identified a total of 10,379 genes that exhibited decreased mH3.3 enrichment in mH3.3-depleted embryos. Using a threshold (fold change >1.2) to pinpoint promoters with significant reductions in H3K27ac after mH3.3 depletion, 77% of these genes (n=8,060) exhibited significantly reduced H3K27ac signals, and thus were classified as mH3.3K27ac-associated genes (**Fig. 4E and S3B**). Notably, within this classification, 3,448 genes had previously been categorized as minor ZGA genes (3), constituting 87% of the genes exhibiting prominent mH3.3K27ac enrichment at promoters in zygotes (**Fig. 4F**). Intriguingly, 1,160 out of 1,275 downregulated minor ZGA genes (90%) in mH3.3-depleted embryos were marked by H3K27ac (**Fig. 4G**). Therefore, these 1,160 genes were designated as mH3.3K27ac-associated minor ZGA genes.

The minor wave of ZGA occurs at the late one-cell and early two-cell stage and is believed to be predominantly driven by the paternal genome (42, 43). This notion is further substantiated by a recent study demonstrating that the paternal genome exhibits more extensive H3K27ac enrichments compared to the maternal genome in zygotes (40). Consistent with these findings, we observed that the enrichments of H3K27ac in the paternal genome were in strong correlation with the patterns of mH3.3 on the paternal genome (**Fig. 4H**). In stark contrast, the maternal genome appeared to be relatively depleted of the H3K27ac modification at the one-cell/zygote stage (**Fig. 4H**). Furthermore, in correlation with the enrichment of mH3.3K27ac on minor ZGA genes, we found that mH3.3K27ac in the paternal genome encompasses 83% (n=929) of the mH3.3K27ac-associated minor ZGA genes, including *Dux* (also known as *Duxf3* in mice), which can activate ERVL-family repeats and ERVL-linked genes during the major ZGA at the late 2-cell stage (44) (**Fig. 4I-J**). In conclusion, our findings suggest that the deposition of mH3.3 is essential for establishing H3K27ac, which in turn is crucial for the activation of minor ZGA genes. This activation is predominantly a consequence of the epigenetic reprogramming of the paternal genome following fertilization.

### Failed mH3.3S31 Phosphorylation Leads to mH3.3K27ac Loss, Minor ZGA Failure and Developmental Arrest

Recent studies have shown that the phosphorylation of histone H3.3 at serine 31 influences the acetylation of adjacent lysine residues, which in turn stimulates induced transcription (38, 45). We thus hypothesize that the phosphorylation of mH3.3 at serine 31 (mH3.3S31p) could be instrumental in establishing mH3.3K27ac, which is essential for the activation of minor ZGA genes. Notably, a strong H3.3S31 phosphorylation (H3.3S31p) signal can be detected in the pronuclei shortly after fertilization (**Fig. 5A**). In alignment with this, we noted a decline in the global levels of H3K27ac signals within the pronuclei of zygotes upon the inhibition of H3.3S31p using SB-218078 (**Fig. S4A, B**), a Checkpoint Kinase 1 (Chk1) inhibitor known to affect H3.3S31p (46). This finding suggests a correlation, whereby H3.3K27ac levels are modulated by H3.3S31p.

**Figure 5.**
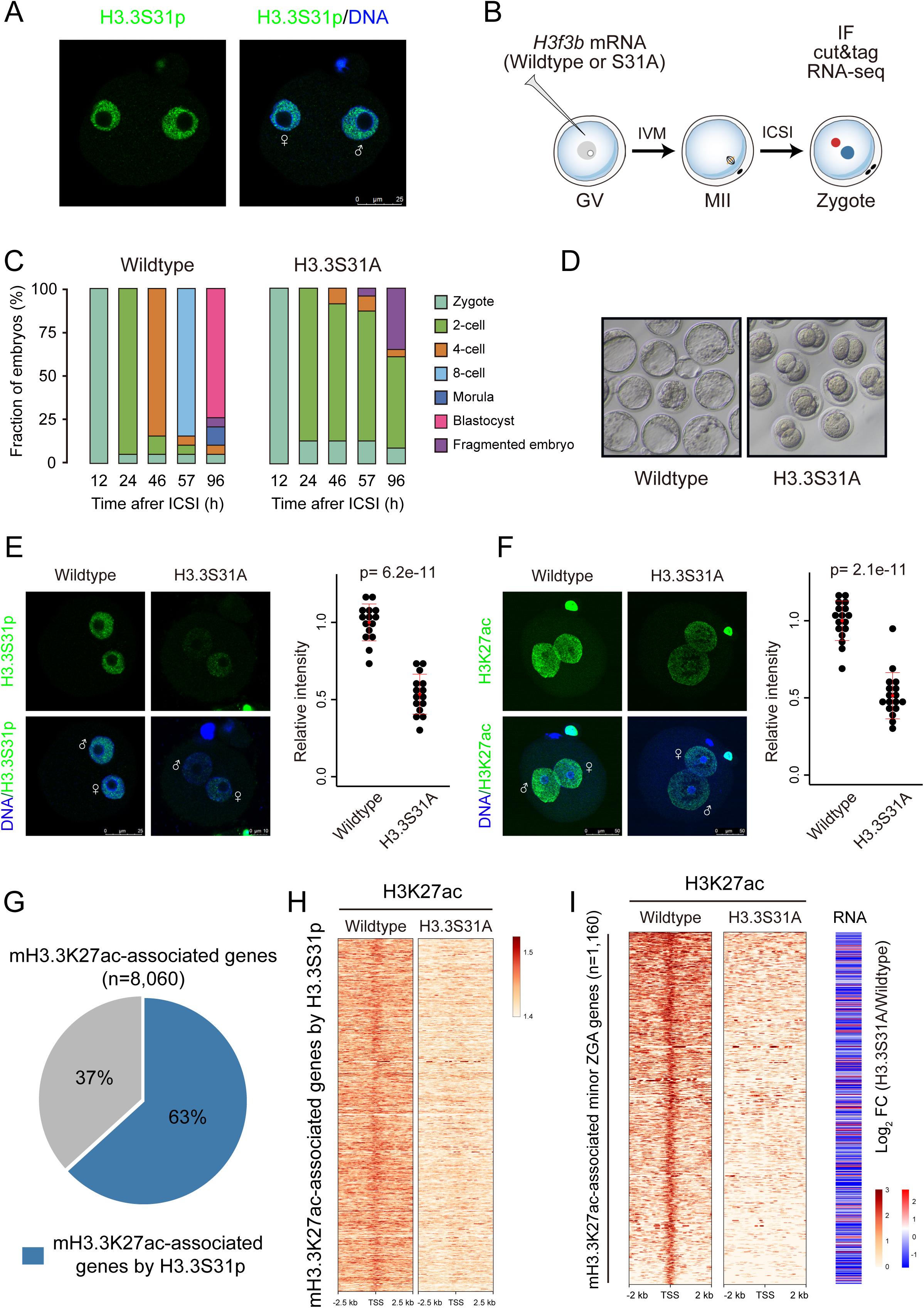
Failed mH3.3S31 phosphorylation leads to mH3.3K27ac loss. (A) Immunostaining for H3.3S31p in zygotes. (B) Schematic presentation of the wildtype H3.3 or H3.3S31A injection experimental protocol. (C) Stacked bar plots showing fraction of mouse embryos at the different developmental stages after wildtype H3.3 or H3.3S31A overexpression. (D) Representative images of the development of wildtype H3.3 or H3.3S31A-overexpressed mouse embryos. Immunofluorescence and fluorescence intensity of H3.3S31p (E) and H3K27ac (F) in wildtype H3.3 or H3.3S31A-overexpressed zygotes. (G) Pie chart showing the fraction of mH3.3K27ac-associated genes lost H3K27ac in H3.3S31A-overexpressed group. (H) Heatmap showing the loss of H3K27ac signals of mH3.3K27ac-associated genes (n=5,066) in H3.3S31A-overexpressed group. (I) Heatmap (left) showing the loss of H3K27ac signals of mH3.3K27ac-associated minor genes (n=1,159) in H3.3S31A-overexpressed group. Heatmap (right) showing the change in expressions of mH3.3K27ac-associated minor ZGA genes (n=1,159). Mean values of two biological replicates are scaled and represented as Z scores.

To further investigate the role of mH3.3 phosphorylation in establishing mH3.3K27ac, we conducted an experiment in which we injected germinal vesicle (GV) oocytes with either wild-type H3.3 mRNA or H3.3S31A mRNA (a non-phosphorylatable mutant). Following the injections, the oocytes underwent in vitro maturation (IVM) and ICSI, and we determined their effect on mH3.3S31p and mH3.3K27ac levels (**Fig. 5B**). The injection of H3.3S31A mRNA had no effect on IVM, as evidenced by the extrusion of the first polar body (**Fig. S4C, D).** H3.3S31A-injected embryos failed to develop beyond the 2-cell stage, a phenotype that mirrors what was observed in mH3.3-depleted embryos (**Fig. 5C, D**). Coinciding with this developmental arrest, we observed a reduction in the levels of H3.3S31p and H3.3K27ac compared to the control group (**Fig. 5E, F**). The substantial reductions in H3K27ac signals were further confirmed in the H3.3S31A-overexpressed group (**Fig. S4E**). These results reveal the importance of mH3.3S31p on early embryonic development and mH3.3K27ac establishment. Notably, approximately 63% of mH3.3K27ac-associated genes exhibited a loss of H3K27ac in the H3.3S31A-overexpressed group (**Fig. 5G, H**). We further analyzed whether embryos overexpressing H3.3S31A exhibited defective minor ZGA progression by employing RNA-seq. Transcriptomic analysis revealed significantly reduced expressions of mH3.3K27ac-associated minor ZGA genes (n=1,160) in the H3.3S31A-overexpressed group (**Figs. 5I and S4F, G**). Collectively, our data suggest a critical role of mH3.3S31p-mediated acetylation at lysine 27 for minor ZGA gene activation, which appears to be vital for early embryonic development.

### mH3.3 Deposition: Reducing Repressive Modifications and Enabling the Onset of Minor ZGA

We found that sH3.3 on the sperm genome may not be completely lost after fertilization. Intriguingly, about half of this sH3.3 enriched loci in the sperm genome is also observed with a high level of mH3.3 enrichment in the androgenetic embryos (**Fig. 6A**). This suggests that after fertilization, this sH3.3 is replaced by mH3.3. Furthermore, the mH3.3K27ac-associated minor ZGA genes (n=1,160) show a higher enrichment of pan-H3 at TSSs compared to both non-mH3.3-dependent ZGA genes and the average level of TSSs across the sperm genome (**Fig. 6B**). This implies that these genes are associated with nucleosomes retained in the sperm genome, as opposed to being packaged by protamines. Notably, these genes are predominantly enriched with H3K27me3 (n=743) in the sperm genome (**Fig. 6C**).

**Figure 6.**
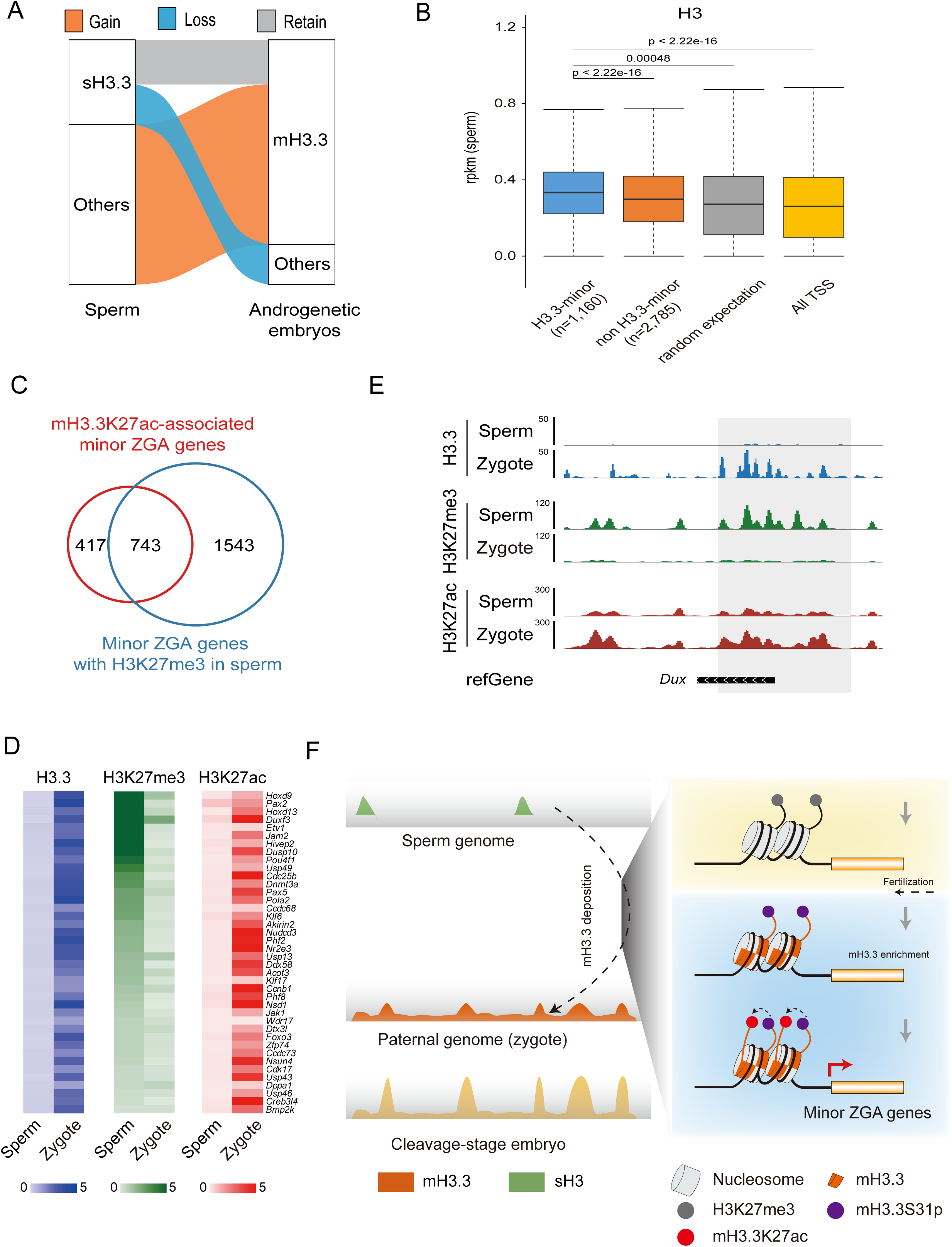
Deposition of mH3.3 on the sperm genome facilitates minor ZGA. (A) The dynamic changes of H3.3 from sperm to embryos. (B) Box plot displaying the enrichment of H3 on the sperm genome for mH3.3-dependent minor ZGA genes (n=1,160) comparing to non H3.3-dependent minor ZGA genes (n=2,785), randomly selected genes and all TSSs. (C) Venn diagrams showing the overlap of mH3.3K27ac-associated minor ZGA genes and minor ZGA genes with H3K27me3 in sperm. (D) Heat maps depicting the enrichment of H3.3, H3K27ac, and H3K27me3 for a selection of minor ZGA genes in sperm and zygotes. (E) Genome browser view illustrating the signals of H3.3, H3K27ac, and H3K27me3 at the *Dux* locus in sperm and zygotes. (F) A model summarizing the landscapes of parental allele-derived H3.3 in mouse sperm genome and cleavage-stage embryos (left). Post-fertilization, mH3.3 significantly aids in the paternal epigenetic remodeling by establishing serine phosphorylation-mediated mH3.3K27ac, facilitating the removal of H3K27me3 present in gametes, and activating minor ZGA (right).

Next, we explored how these modifications evolve following fertilization and during ZGA. Focusing on the mH3.3K27ac-associated minor ZGA genes, we observed a significant increase in H3.3 deposition coupled with a decrease in H3K27me3 modifications at TSSs, concurrently, there was an increase in active H3K27ac modification in the zygote throughout ZGA (**Figs. 6D**). Intriguingly, the 743 minor ZGA genes, which initially exhibited low H3.3 enrichment and high H3K27me3 modifications in the sperm genome, transitioned to greater H3.3 enrichment and reduced H3K27me3 modification in the zygote. This transition also entailed the establishment of H3K27ac during ZGA, exemplified by the *Dux* gene (**Fig. 6E**). These observations suggest that the deposition of mH3.3 is inversely correlated with the loss of repressive modification H3K27me3, while being positively related to the increase of active H3K27ac modification during ZGA. Based on these findings, we propose that the deposition of mH3.3 displaces the repressive histone modifications, such as H3K27me3, in the sperm genome, and creates a permissive chromatin state that is conducive to establishing the active H3K27ac modification, thus enabling the initiation of minor ZGA (**Fig. 6F**).

## Discussion

In our study using H3.3B-HA reporter mice, we explored the differential deposition of maternally and paternally expressed H3.3 on the zygotic genome during early embryogenesis. We observed that H3.3 (H3.3B) from the paternal allele begins its deposition during minor ZGA at the early two-cell stage. In contrast, the deposition of H3.3 from the maternal allele is delayed until the four-cell stage. Given these observations, we further examined the epigenetic processes behind paternal genome reprogramming during fertilization and ZGA. We discovered that maternal H3.3 (mH3.3) plays a crucial role in reprogramming the paternal genome and activating minor ZGA genes. Mechanistically, mH3.3 deposition on the paternal genome starts globally with a non-canonical pattern, as described in a previous study on mature oocytes using H3.3 antibody (22). However, it shows preferential enrichment at CpG-rich TSSs following the protamine-to-histone transition and the first embryonic division. This deposition of mH3.3 on the paternal genome is closely associated with minor ZGA gene activation and the establishment of the H3K27ac modification. Additionally, a failure in H3S31 phosphorylation leads to a loss of H3K27ac modification, resulting in minor ZGA disruption and subsequent developmental arrest. Our research highlights the pivotal role of mH3.3, an essential maternal H3 histone variant, in paternal genome reprogramming and the onset of minor ZGA. This process is facilitated through H3.3 replacement, a mechanism that not only eliminates repressive histone modifications on the sperm genome but also simultaneously introduces new sites for histone modifications that promote gene activation.

In mice, zygotic transcription is initiated shortly after pronucleus formation in 1-cell embryos. Yet, the specific transcriptional loci and the regulatory mechanisms behind their expression are not fully understood (47). Although the majority of minor ZGA transcripts are thought to be promiscuously expressed across diverse genomic regions, including intergenic areas (47–49), our research identifies 1,160 mH3.3K27ac-associated minor ZGA genes that are regulated by mH3.3. Notably, 743 of these genes including *h3f3b* gene display H3K27me3 enrichment at the TSS in the sperm genome. This number closely aligns with the 817 genes specifically identified as being expressed in fertilized embryos, presumably from the paternal genome (3). Our results underscore that the activation of these minor ZGA genes requires mH3.3 deposition at their TSSs, a process enhanced by the establishment of mH3.3K27ac and mH3.3S31p modifications in promoter regions. Therefore, our findings indicate that while many previously identified minor ZGA genes appear to be promiscuously expressed, 743 of them including *h3f3b* gene are specifically regulated by mH3.3 and are expressed from the paternal genome rather than from the maternal genome during the process of minor ZGA.

The epigenetic landscapes of the sperm and oocyte genomes are distinct, with each undergoing its own unique reprogramming process post-fertilization. Although the paternal genome has been shown to activate at the late one-cell stage, the activation timing of the maternal genome remains unclear. A key challenge in studying this issue is to distinguish between pre-existing maternal RNAs and the newly transcribed ones from the maternal genome after fertilization. This differentiation is particularly challenging with RNA-seq technology since the maternal stored RNAs are considerably more abundant than the de novo transcripts from the maternal genome in the 1- and 2-cell stage embryo. In our study, using H3.3B-HA reporter mice, we discovered that H3.3 (H3.3B-HA) expressed from both paternal and maternal genomes have similar distribution profiles from the 2-cell stage onward, which is specifically enriched at the CpG-rich TSSs, contrasting to the evenly distribution across the whole genome in the oocyte and zygote. Notably, we observed a significant increase in the deposition of H3.3B-HA expressed from the paternal allele on the zygotic genome starting at the 2-cell stage, while the deposition of H3.3B-HA from the maternal allele was delayed until the 4-cell stage. This suggests that the activation of paternal genome is in the first wave of ZGA, whereas the activation of maternal genome is delayed in the second wave of ZGA. However, the detailed mechanism of maternal genome activation warrants further investigation.

In our studies with androgenetic embryos, we identified a dual role for mH3.3 in the reprogramming of the paternal genome. Initially, mH3.3 acts as the primary H3 variant, facilitating the formation of nucleosomes across the paternal genome in a non-canonical pattern during the protamine-to-histone transition prior to DNA replication. Subsequently, mH3.3 becomes selectively enriched at CpG-rich promoters. The selective enrichment we observed might correlate with the increased turnover of H3.3 at the CpG-rich loci (**Fig. 2F, S2G-H**). These results support a CpG-associated but functionally random pattern of early deposition of mH3.3B, complemented by a CpG-associated non-random deposition of mH3.3 near the promoters and TSSs for genes that function in early development. This randomness decreases over time, whereas the CpG-associated pattern increases after the first embryonic DNA replication and remains dominant throughout the preimplantation stage. The mechanism underlying the preferential deposition of mH3.3 and how it is regulated by DNA replication remains to be further elucidated.

Previous studies indicate that the sperm genome’s residual nucleosomes primarily contain specific modifications like H3K4me3 and H3K27me3. These unique nucleosome patterns at gene promoters are believed to act as epigenetic markers, potentially priming the ZGA genes for post-fertilization activation. Our findings show that among the 1,160 mH3.3-associated minor ZGA genes, there’s significant H3 histone retention at their TSS in the sperm genome, predominantly enriched with H3K27me3. This suggests these genes are pre-patterned for activation. H3.3 incorporation into chromatin, especially at gene bodies and promoters, makes chromatin more accessible, aiding in transcriptional activation, especially through histone acetylation which promotes open chromatin. Our data highlight the crucial role of mH3.3S31p-mediated acetylation in minor ZGA gene activation, vital for early embryonic development.

In summary, our research underscores the distinct activation patterns of paternal and maternal genomes, spotlighting a set of mH3.3-regulated minor ZGA genes primarily derived from the paternal genome. These genes are characterized by discernible pre-patterning, marked by nucleosome retention and H3K27me3 enrichment at their TSSs in the sperm genome. Mechanistically, mH3.3 replaces H3, which carries repressive modifications, such as H3K27me3, in the sperm genome. This replacement with de novo synthesized H3.3 eliminates the repressive modifications and enriches H3.3 at the loci, promoting a more permissive chromatin state. Simultaneously, it provides new modification sites for establishing active modifications such as H3K27ac and H3K4me3, leading to minor ZGA initiation.

## Methods

### Animals and oocytes

Animals were housed and prepared in accordance with the protocol approved by the IACUC of Weill Cornell Medical College (Protocol number: 2014-0061). B6D2F1 and ICR mice were obtained from Taconic Farms (Germantown, NY). Female mice, aged 6-8 weeks, were superovulated using 5 IU of PMSG (Pregnant mare serum gonadotrophin, Sigma-Aldrich, St. Louis, MO) and 5 IU of hCG (Human chorionic gonadotrophin, Sigma-Aldrich) with a 48-hour interval between injections. MII oocytes were collected from superovulated female mice 14-16 hours after hCG administration. Embryos were cultured in KSOM medium (MR-121-D; Merck, Kenilworth, NJ, USA) under mineral oil at 37°C in a 5% CO2 incubator.

### Treatment of oocytes with inhibitors and chemicals

SB-218078 (HY-107407; MedChemExpress, Monmouth Junction, NJ, USA) was dissolved in dimethyl sulfoxide (DMSO). This solution was then diluted to achieve the desired concentration in the maturation medium (SB-218078, 3 µM). Embryos were cultured *in vitro* using KSOM medium with varying doses of the inhibitor for subsequent analysis. Additionally, 0.05% DMSO served as a negative control.

### Plasmid construction

cDNAs encoding human *h3f3b* were subcloned into expression plasmids driven by the T7 promoter for in vitro transcription of mCherry-tagged sequences. A non-inhibitable *h3f3b* variant was constructed by replacing the S31 residue of human *h3f3b* with an alanine.

### In vitro transcription

To prepare mRNAs for microinjection, we linearized the constructed vectors and purified them using phenol-chloroform, followed by ethanol precipitation. The linearized DNAs were then transcribed in vitro with the mMESSAGE mMACHINE Kit (AM1344; Invitrogen, Carlsbad, CA, USA). The transcribed mRNAs were polyadenylated, adding poly(A) tails of 200–250 bp, using the mMACHINE Kit (AM1350; Invitrogen). These mRNAs were then recovered via lithium chloride precipitation and resuspended in nuclease-free water. The resulting synthetic mRNA was frozen and stored at −80 °C.

### Microinjection

MII oocytes were harvested from superovulated B6D2F1 or ICR females 14-16 hours post-hCG injection. The microinjection process utilized a FemtoJet 4i microinjector (Eppendorf, Hamburg, Germany) in tandem with ECLIPSE Ti micromanipulators (Nikon, Tokyo, Japan). Both siRNAs and morpholinos were delivered into the oocytes using a piezo-operated microcapillary pipette featuring a 3-5 µm inner diameter. After the injection, the oocytes were left to rest at room temperature for 10 minutes, then shifted to an incubator for another 30 minutes at a minimum. Subsequently, either ICSI or parthenogenetic activation was executed. All siRNAs employed in this research were sourced from RiboBio, while the morpholinos were obtained from GENE TOOLS (Philomath, OR, USA). The specific sequences for H3f3a and H3f3b morpholinos and siRNAs can be found in Supplementary Table 2. For the overexpression of H3.3S31A in GV oocytes, we collected fully grown GV oocytes from PMSG-primed (44 h) 23-day-old mice. To prevent spontaneous GVBD, these GV oocytes were harvested in M2 medium supplemented with 2M milrinone. An amount of approximately 5–10 pL of 250ng/µl mRNAs was microinjected into the oocyte’s ooplasm. These injected oocytes underwent culture in G-1 plus medium containing 2M milrinone at 37℃ with 5% CO2 for 6 hours to facilitate the translation of the introduced mRNAs. Subsequently, the oocytes were moved to milrinone-free G-1 plus medium. The maturation status of the MII oocytes was assessed 14-16 hours after their release from the milrinone treatment.

### Enucleation and Intracytoplasmic Sperm Injection (ICSI)

Oocytes from both H3.3B-HA and B6D2F1 were placed in a droplet of HEPES-CZB medium containing 5 µg/ml cytochalasin B, which was prepositioned in the operation chamber on the microscope stage. Utilizing a micromanipulator and a precision glass needle, the spindle-chromosomal complex (SCC) was meticulously extracted from both the donor H3.3B-HA oocytes and the recipient B6D2F1 oocytes. The SCC from the H3.3B-HA oocyte was subsequently introduced into the perivitelline space of the enucleated B6D2F1 oocyte. These oocytes then underwent electrofusion: they were aligned and subjected to a sequence of electrical pulses to facilitate membrane fusion. After fusion, the oocytes were activated by being cultured in Ca2+-free CZB medium enriched with 10 mM Sr2+ and 5 µg/ml cytochalasin B for a duration of 5 hours. Following this, they were relocated to an incubator and nurtured in advanced KSOM (MR-101-D, Merck) at 37°C with 5% (v/v) CO2 in air.

For ICSI, a WT sperm head from ICR mice was selected in the PVP droplet using the injection pipette, and each H3.3B-HA oocyte received an injection of one sperm head. To explore the dynamics of histone H3.3 deposition in early mouse embryos, embryos maternally expressing H3.3B-HA (where a WT sperm was introduced into an H3.3B-HA oocyte) and those expressing the paternal allele of H3.3B-HA (in which H3.3B-HA sperm was introduced into a WT oocyte) were harvested separately. For androgenetic embryos, two WT sperm heads were introduced into enucleated oocytes sourced from H3.3B-HA reporter mice. Post-ICSI, embryos were permitted to rest at room temperature for about 10 minutes before being moved to KSOM medium and incubated at 37°C with 5% (v/v) CO2 in air.

### ULI-NChIP–seq

For ULI-NChIP–seq, we utilized approximately 100 embryos for each reaction, conducting two to three replicates for every stage. All harvested embryos underwent three wash cycles in a 0.5% BSA in PBS solution to mitigate potential contamination. The ULI-NChIP process was executed as described in a previous study (28). For each immunoprecipitation reaction, we employed one microgram of the HA antibody (9727, ab9134, Abcam, Cambridge, CB2 0AX, UK). Sequence libraries were crafted using the KAPA Hyper Prep Kit, designed for the Illumina platform (kk8504), in adherence to the manufacturer’s guidelines. All the libraries were subsequently sequenced on the Illumina platform, following the specified manufacturer’s protocols.

### CUT&Tag

CUT&Tag was conducted using the Hyperactive In-Situ ChIP Library Prep Kit for Illumina (TD903; Vazyme Biotech, Nanjing, China). For mH3.3-depleted and H3.3S31A-injected zygotes, both H3.3 and H3K27ac CUT&Tag procedures were executed 10-12h post-ICSI. Embryos were treated with 10 μl of pre-washed ConA beads. To this, 50 μl of antibody buffer and 0.5 μg of the primary antibody were added. The solution was then left to incubate overnight at 4°C. After two washes using dig-wash buffer, the embryos were incubated in 150 μl of the same buffer, now containing 0.7 μg of the secondary antibody, for an hour at room temperature. Following another pair of washes with 800 μl dig-wash buffer, we added 0.3 μl of pG-Tn5 and 50 μl of dig-300 buffer to the samples and incubated them for another hour at room temperature. This was followed by two more washes with 800 μl of dig-wash buffer. Subsequently, 300 μl of tagmentation buffer was added, and the samples were incubated at 37°C for an hour. To terminate the reaction, we added 10 μl of 0.5 M EDTA, 3 μl of 10% SDS, and 2.5 μl of 20 mg/ml proteinase K. The samples then underwent phenol-chloroform extraction and ethanol precipitation. PCR amplification of the libraries was carried out under these conditions: an initial step at 72°C for 5 minutes, denaturation at 98°C for 30 seconds, followed by 20 cycles of 98°C for 10 seconds, and annealing at 60°C for 30 seconds. This was concluded with an extension step at 72°C for 1 minute before holding the samples at 4°C. The subsequent clean-up post-PCR involved the addition of 1.5× volume of DNA Clean Beads (N411; Vazyme Biotech). Libraries were allowed to interact with the beads for 10 minutes at room temperature. After a gentle rinse with 80% ethanol, the libraries were eluted in 20 μl of elution buffer. Sequencing of all libraries was performed on the Illumina NovaSeq 6000 platform, adhering to the manufacturer’s guidelines.

### ChIP-seq and CUT&Tag data processing

ChIP-seq and CUT&Tag reads were aligned to the mouse genome (mm10) using Bowtie 2 (version 2.2.5) with a default setting after removing adaptor sequences and low-quality reads by fastp (version 0.12.4). Data reproducibility between replicates was assessed by correlation analysis of mapped read counts across the genome. Then, we pooled the biological replicates for each stage and performed downstream analysis. Drosophila genome DNA was used as spike-in in our experiment. When spike-in was included, reads were mapped to the reference genome in which mm10 and Drosophila genome (Drosophila_melanogaster.BDGP6.32.dna.toplevel) are combined. Reads from PCR duplicates were removed using Picard. After confirming reproducibility between replicates, they were merged using Samtools. The mouse genome (mm10) was divided into 10 kb bins in a sliding window of 5 kb. The number of ChIP and input reads covering each bin was calculated using the BEDtools intersect with option ‘-c’. To compute normalized enrichment, normalized ChIP read counts were divided by normalized input read counts. Final bam files were converted to bigWig files of read coverages normalized to RPKM using deepTools (version 3.5.1) bamCoverage. Heatmaps were depicted using computeMatrix and plotHeatmap in deepTools (version 3.5.1) for genes. Genomic regions (10 kb bin) with ≥0.2 normalized H3.3 enrichment were considered as H3.3-enriched regions. Genomic regions (10 kb bin) with ≥0.1 normalized H3 enrichment were considered as H3-enriched regions. The washU epigenome browser (http://epigenomegateway.wustl.edu/browser/) was used to visualize ChIP-seq and CUT&Tag data.

### Histone modification analyses

H3K27ac (GSE207222)(55), H3K36me3 (GSE112834)(57), H3K4me3 and H3K27me3 (GSE73952)(28), H3K9me3 (GSE98149)(58) and H3-sperm (GSE79227)(59) ChIP-seq datasets were downloaded from previous publications. ChIP-seq reads were aligned to the mouse genome (mm10) using Bowtie 2 after removing adaptor sequences and low-quality reads using Fastp. Duplicate and low-mapping-quality (MAPQ <5) reads were removed. Heatmaps were generated using deepTools. Final bam files were converted to bigWig files of read coverages normalized to RPKM using deepTools (version 3.5.1) bamCoverage. Heatmaps were depicted using computeMatrix and plotHeatmap in deepTools (version 3.5.1) for autosomal gene. CG site (observed versus expected) was calculated for ±2.5 kb from TSSs.

### Identification of aberrant and ZGA gene associated H3K27ac domains

For the zygote stages, we quantified the expression levels for each putative H3K27ac domain using the deeptools tool and compared H3K27ac levels between the control and experimental groups using the DESeq2 package in R. We identified H3K27ac-enriched regions by Macs2 callpeaks with the option ‘-p 0.05 –nomodel’. And the aberrant H3K27ac domains of the stage were identified using stringent criteria: sum of the normalized H3K27ac levels of the control and treated groups ≥ 1.2 in H3K27ac-enriched regions.

The predefined ZGA genes list was from the previous study(3). Promoters of the ZGA genes (defined as 1 kb around the transcription start site) that overlapped with the aberrant H3K27ac domains according to the BEDTools intersect tool were defined as ZGA gene H3K27ac domains. Changes in CUT&Tag data were processed and visualized using the computeMatrix and plotHeatmap functions of deepTools. Data were imported using the default settings and all values were normalized to RPKM values and scaled as described above. The retained regions were merged using the merge function in BEDTools v2.30.0 if they were consecutive or overlapped.

### RNA-seq

RNA-seq libraries were prepared as previously described(54). For mH3.3-depleted and H3.3S31A-injected zygotes, RNA-seq were performed at 10-12h post-ICSI. Briefly, three embryos were used per reaction, and two replicates were performed for each group. All embryos were washed three times in 0.5% BSA-PBS solution to avoid possible contamination. DNA was amplified using the Phusion Hot Start II High-Fidelity PCR Master Mix (F-565S; Thermo Fisher). Library preparation was performed using the TruePrep DNA Library Prep Kit (TD501; Vazyme Biotech, Nanjing, China) according to the manufacturer’s instructions. All libraries were sequenced using the NovaSeq 6000 (Illumina, San Diego, CA, USA) according to the manufacturer’s instructions.

### RNA-seq data processing

For RNA-seq analysis of early stage embryos, FastQC was performed for Illumina reads. We used the Trim Galore software to discard low-quality reads, trim adaptor sequences, and eliminate poor-quality bases. Then, we downloaded the mouse reference genome (genome assembly: mm10) from the Ensembl database and used the HISAT2 software for read alignment. The gene-level quantification approach was used to aggregate raw counts of mapped reads using the featureCounts tool. The expression level of each gene was quantified in terms of the normalized fragments per kilobase of transcript per million mapped reads (FPKM). Next, we used the R package DESeq2 for differential gene expression analysis. KEGG analysis of screened DEGs was performed using the KOBAS online tool (http://kobas.cbi.pku.edu.cn/kobas3/).

### Immunofluorescence staining

Mouse embryos were fixed in fixative solution (FB002; Invitrogen) for 40 min at room temperature, followed by permeabilization in 1% Triton X-100 (93443, 100 ml; Sigma) for 20 min at room temperature. Embryos were then blocked in blocking solution consisting of 1% bovine serum albumin (BSA) in phosphate-buffered saline (PBS) for 1 h at room temperature after three washes in washing solution (0.1% Tween-20, 0.01% Triton X-100 in PBS). Antibody incubation (HA, ab9134, Abcam, Cambridge, CB2 0AX, UK; Histone H3.3, 91191, Active Motif, Carlsbad, CA, USA; H3.3S31p, ab92628, Abcam, Cambridge, CB2 0AX, UK; H3K27ac: 39034; Active Motif, Carlsbad, CA, USA) was performed overnight at 4°C. The next day, the embryos were washed in washing solution and incubated with secondary antibodies (Invitrogen) for 1 h at room temperature. After staining with Hoechst, the embryos were washed in washing solution. Embryo imaging was performed using an inverted confocal microscope (TCS SP8; Leica, Wetzlar, Germany) and analyzed using LAS X software (Leica).

### Statistical analyses

All statistical analyses were performed using R v4.1.0 software (R Development Core Team, Vienna, Austria). Data are expressed as means ± standard error of the mean (SEM). Differences between means were evaluated using the two-tailed Student’s t-test or Wilcoxon rank sum test. Asterisks indicate significant differences as follows: *P < 0.05, **P < 0.01, and ***P < 0.001.

### Data Availability, Code Availability

RNA-Seq, ChIP-Seq and CUT&Tag data that support the findings of this study have been deposited in the GEO under accession code GSE242960. Previously published ChIP-Seq data that were re-analyzed here are available under accession codes GSE73952 (H3K4me3 and H3K27me3), GSE207222 (H3K27ac), GSE79227 (H3-sperm). All other data supporting the findings of this study are available from the corresponding author upon reasonable request.

---

**Figure S1.**
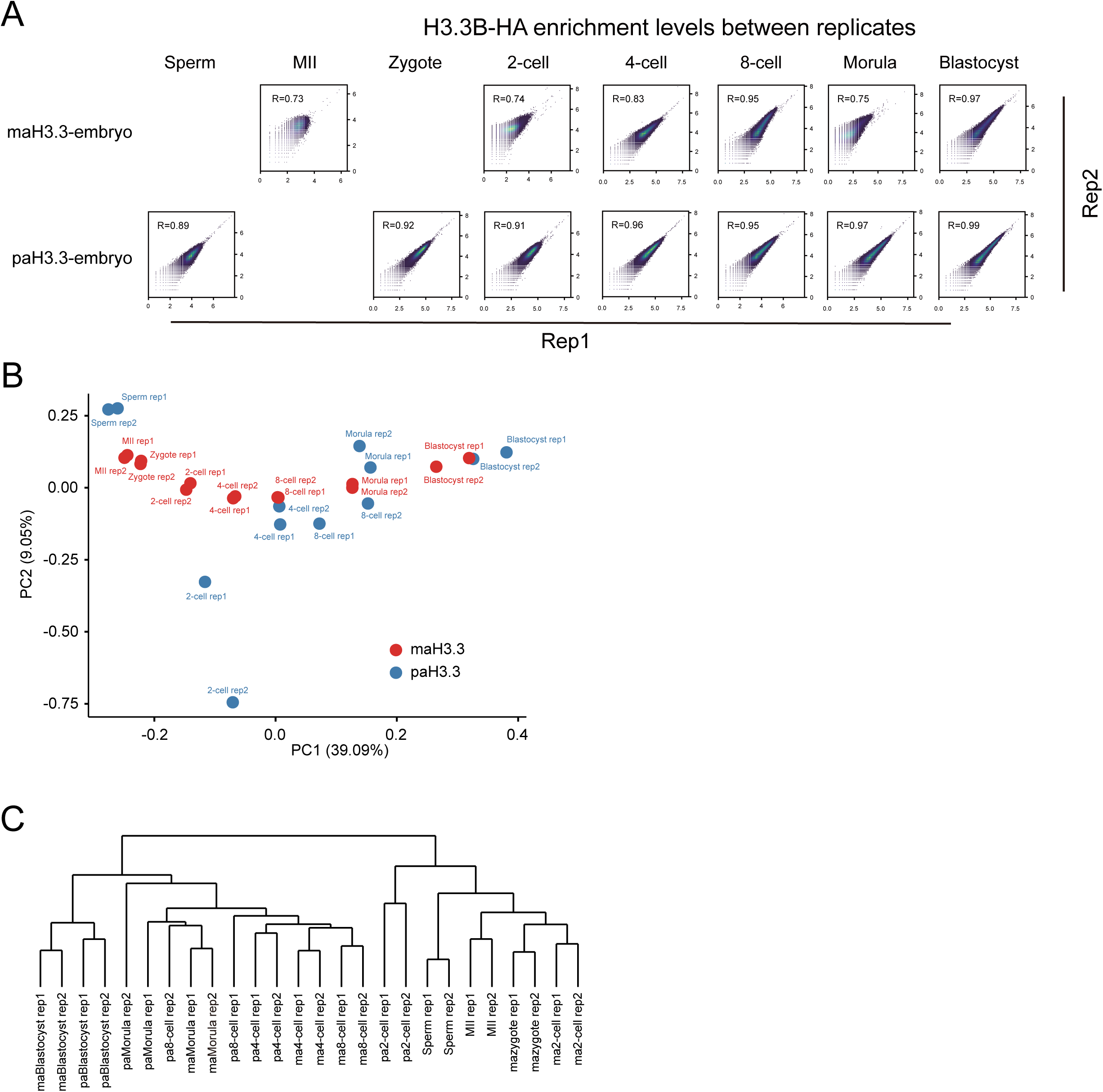
Validation of H3.3 ULI-NChIP. (A) Correlation analysis between replicates within each group. Pearson correlation coefficients were calculated using RPKM (Reads Per Kilobase Million) values across 10-kb-binned regions encompassing the entire genome. (B) Hierarchical clustering of global maH3.3 and paH3.3 enrichments in different samples. (C) Principal Component Analysis (PCA) executed based on H3.3 enrichment in mouse gametes and pre-implantation embryos.

**Figure S2.**
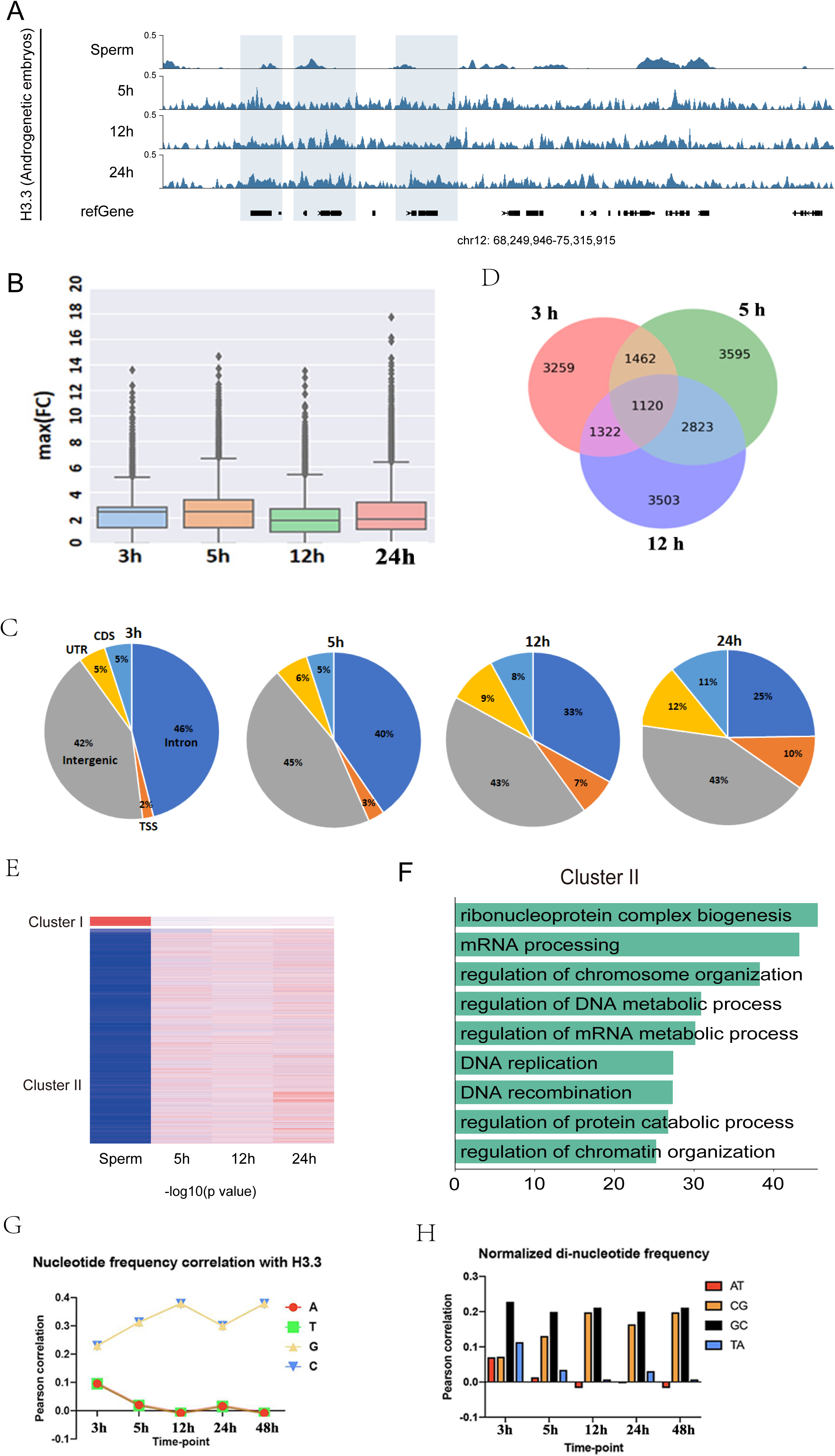
Deposition of mH3.3 onto the Paternal Genome Post-Fertilization. (A) A graph displaying the changes in mH3.3 coverage across various genomic annotations at different time points following fertilization. (B) Distribution of maximum fold change (indicating the presence of H3.3 enrichment) at each time point. (C) A graph displaying the changes in mH3.3 coverage across various genomic annotations at different time points following fertilization. (D) An illustration of the temporal changes in genes enriched for mH3.3 in their TSS. The Venn diagram shows the number of genes that have overlapping replicated H3.3 peaks at various time points. (E) Dynamic changes of mH3.3 patterns in androgenetic embryos. All the mH3.3 domains from sperm to 2-cell were classified into two major clusters using k-means clustering. (F) KEGG analysis of cluster II genes. (G) Relative composition of A, T, G, and C within H3.3 replicated broad peaks after replication. (H) Correlation of H3.3 broad peaks with CpG dinucleotides.

**Figure S3.**
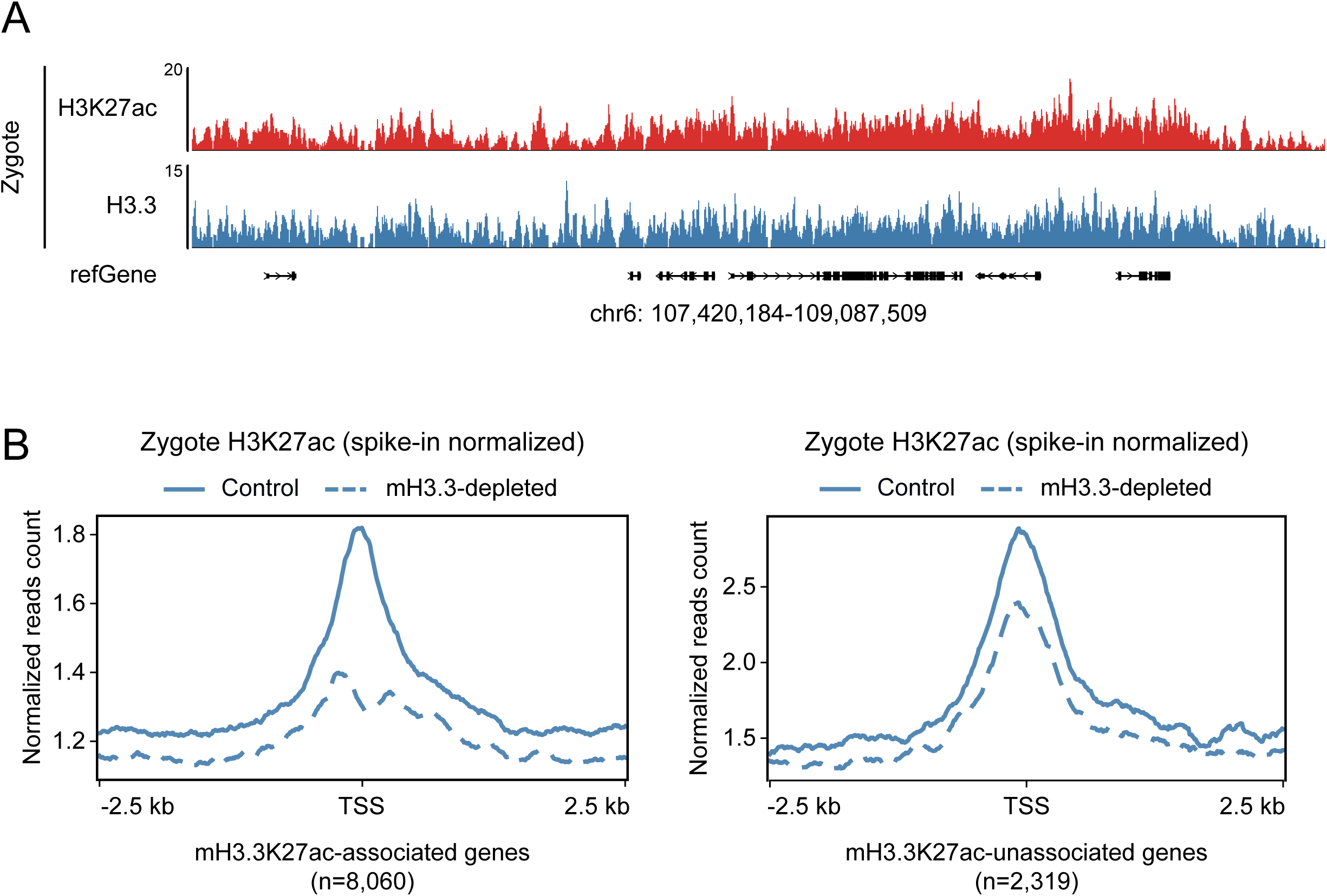
Correlation of H3.3 and H3K27ac enrichments in zygote. (A) Genome browser view demonstrating the parallel enrichment patterns of H3.3 and H3K27ac signals during the zygote stage. (B) Metaplot showcasing H3K27ac signals (normalized using spike-in and Z-score) at mH3.3K27ac-associated genes (n=8,060) compared to mH3.3K27ac-unassociated genes (n=2,319).

**Figure S4.**
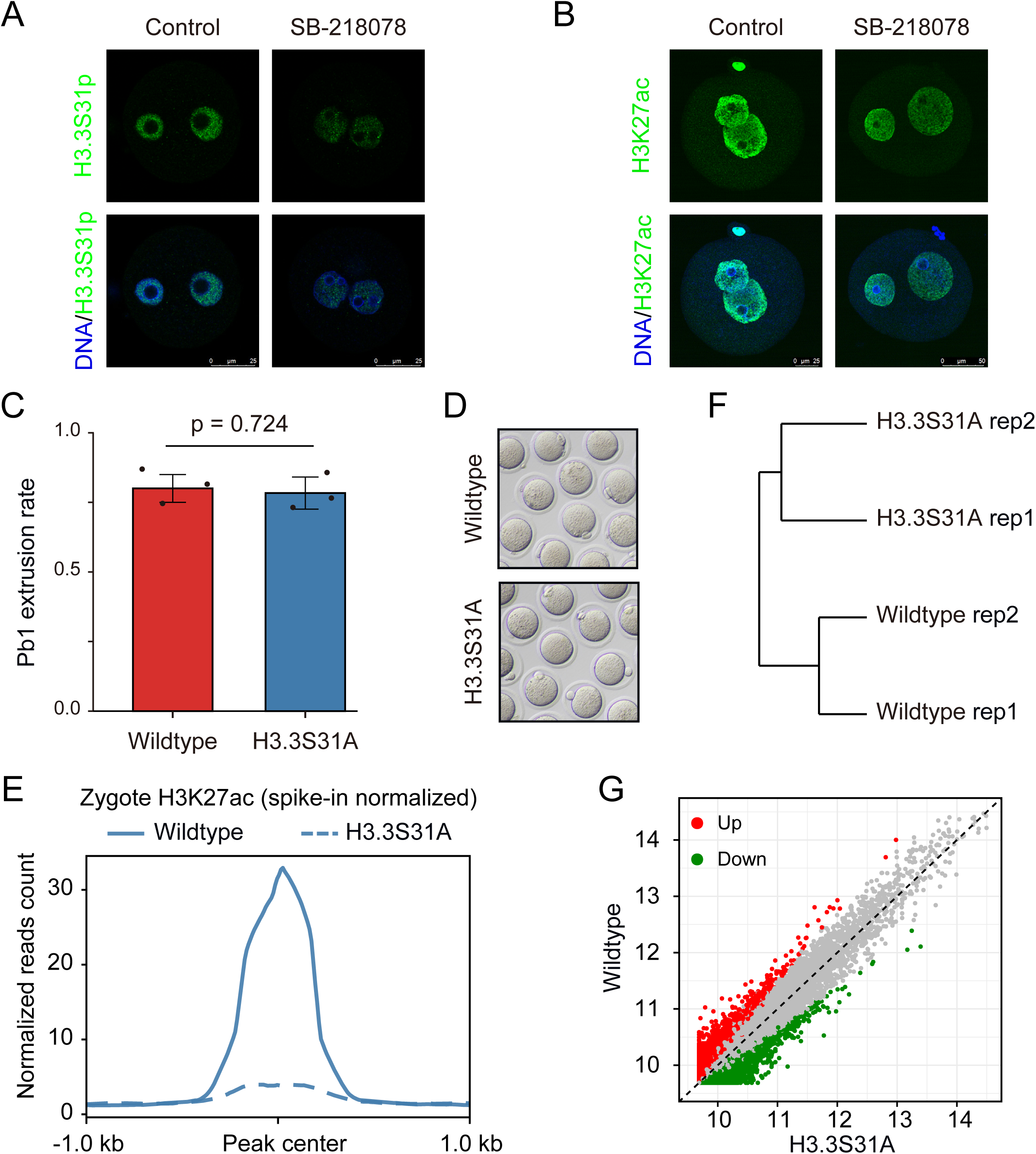
Transcriptomic Analysis of Mouse Embryos Overexpressing Wild-Type H3.3 or H3.3S31A. (A) Immunofluorescence depicting H3.3S31p levels in SB-218078-treated embryos. (B) Immunofluorescence showing H3K27ac levels in SB-218078-treated embryos. (C) Graphical representation indicating that the injection of H3.3S31A mRNA does not affect the in vitro maturation (IVM) of mouse oocytes. (D) Representative images of in vitro matured oocytes following injection of either wild-type H3.3 or H3.3S31A. (E) Metaplot displaying H3K27ac signals (spike-in normalized, Z-score normalized) in mouse embryos overexpressing either wild-type H3.3 or H3.3S31A. (F) Unsupervised clustering analysis illustrating gene expression patterns among mouse embryos overexpressing either wild-type H3.3 or H3.3S31A. (G) Scatter plots depicting the changes in the transcriptome of mouse embryos overexpressing either wild-type H3.3 or H3.3S31A.

## References

1. Clift, D., and Schuh, M. (2013) Restarting life: fertilization and the transition from meiosis to mitosis. Nature reviews. Molecular cell biology 14, 549–562

2. Lee, M. T., Bonneau, A. R., and Giraldez, A. J. (2014) Zygotic Genome Activation During the Maternal-to-Zygotic Transition. Annu Rev Cell Dev Bi 30, 581–613

3. Park, S. J., Komata, M., Inoue, F., Yamada, K., Nakai, K., Ohsugi, M., and Shirahige, K. (2013) Inferring the choreography of parental genomes during fertilization from ultralarge-scale whole-transcriptome analysis. Genes & development 27, 2736–2748

4. Xue, Z. G., Huang, K., Cai, C. C., Cai, L. B., Jiang, C. Y., Feng, Y., Liu, Z. S., Zeng, Q., Cheng, L. M., Sun, Y. E., Liu, J. Y., Horvath, S., and Fan, G. P. (2013) Genetic programs in human and mouse early embryos revealed by single-cell RNA sequencing. Nature 500, 593-+

5. Abe, K., Funaya, S., Tsukioka, D., Kawamura, M., Suzuki, Y., Suzuki, M. G., Schultz, R. M., and Aoki, F. (2018) Minor zygotic gene activation is essential for mouse preimplantation development. Proceedings of the National Academy of Sciences of the United States of America 115, E6780–E6788

6. Dahl, J. A., Jung, I., Aanes, H., Greggains, G. D., Manaf, A., Lerdrup, M., Li, G. Q., Kuan, S., Li, B., Lee, A. Y., Preissl, S., Jermstad, I., Haugen, M. H., Suganthan, R., Bjoras, M., Hansen, K., Dalen, K. T., Fedorcsak, P., Ren, B., and Klungland, A. (2016) Broad histone H3K4me3 domains in mouse oocytes modulate maternal-to-zygotic transition. Nature 537, 548-+

7. Zhang, B. J., Zheng, H., Huang, B., Li, W. Z., Xiang, Y. L., Peng, X., Ming, J., Wu, X. T., Zhang, Y., Xu, Q. H., Liu, W. Q., Kou, X. C., Zhao, Y. H., He, W. T., Li, C., Chen, B., Li, Y. Y., Wang, Q. J., Ma, J., Yin, Q. Z., Kee, K., Meng, A. M., Gao, S. R., Xu, F., Na, J., and Xie, W. (2016) Allelic reprogramming of the histone modification H3K4me3 in early mammalian development. Nature 537, 553-+

8. Schulz, K. N., and Harrison, M. M. (2019) Mechanisms regulating zygotic genome activation. Nat Rev Genet 20, 221–234

9. Eckersley-Maslin, M. A., Alda-Catalinas, C., and Reik, W. (2018) Dynamics of the epigenetic landscape during the maternal-to-zygotic transition. Nature reviews. Molecular cell biology 19, 436–450

10. Ladstatter, S., and Tachibana, K. (2019) Genomic insights into chromatin reprogramming to totipotency in embryos. J Cell Biol 218, 70–82

11. Guo, H. S., Zhu, P., Yan, L. Y., Li, R., Hu, B. Q., Lian, Y., Yan, J., Ren, X. L., Lin, S. L., Li, J. S., Jin, X. H., Shi, X. D., Liu, P., Wang, X. Y., Wang, W., Wei, Y., Li, X. L., Guo, F., Wu, X. L., Fan, X. Y., Yong, J., Wen, L., Xie, S. X., Tang, F. C., and Qiao, J. (2014) The DNA methylation landscape of human early embryos. Nature 511, 606-+

12. Smith, Z. D., and Meissner, A. (2013) DNA methylation: roles in mammalian development. Nat Rev Genet 14, 204–220

13. Amouroux, R., Nashun, B., Shirane, K., Nakagawa, S., Hill, P. W. S., D’Souza, Z., Nakayama, M., Matsuda, M., Turp, A., Ndjetehe, E., Encheva, V., Kudo, N. R., Koseki, H., Sasaki, H., and Hajkova, P. (2016) De novo DNA methylation drives 5hmC accumulation in mouse zygotes. Nat Cell Biol 18, 225-+

14. Hammoud, S. S., Nix, D. A., Zhang, H. Y., Purwar, J., Carrell, D. T., and Cairns, B. R. (2009) Distinctive chromatin in human sperm packages genes for embryo development. Nature 460, 473–U447

15. Torres-Padilla, M. E., Bannister, A. J., Hurd, P. J., Kouzarides, T., and Zernicka-Goetz, M. (2006) Dynamic distribution of the replacement histone variant H3.3 in the mouse oocyte and preimplantation embryos. Int J Dev Biol 50, 455–461

16. Carrell, D. T., and Hammoud, S. S. (2010) The human sperm epigenome and its potential role in embryonic development. Mol Hum Reprod 16, 37–47

17. Frank, D., Doenecke, D., and Albig, W. (2003) Differential expression of human replacement and cell cycle dependent H3 histone genes. Gene 312, 135–143

18. Wellman, S. E., Casano, P. J., Pilch, D. R., Marzluff, W. F., and Sittman, D. B. (1987) Characterization of mouse H3.3-like histone genes. Gene 59, 29–39

19. Wen, D. C., Banaszynski, L. A., Liu, Y., Geng, F. Q., Noh, K. M., Xiang, J., Elemento, O., Rosenwaks, Z., Allis, C. D., and Rafii, S. (2014) Histone variant H3.3 is an essential maternal factor for oocyte reprogramming. Proceedings of the National Academy of Sciences of the United States of America 111, 7325–7330

20. Guo, P. P., Liu, Y., Geng, F. Q., Daman, A. W., Liu, X. Y., Zhong, L. W., Ravishankar, A., Lis, R., Duran, J. G. B., Itkin, T., Tang, F. Y., Zhang, T., Xiang, J., Shido, K., Ding, B. S., Wen, D. C., Josefowicz, S. Z., and Rafii, S. (2022) Histone variant H3.3 maintains adult haematopoietic stem cell homeostasis by enforcing chromatin adaptability (vol 24, pg 99, 2022). Nat Cell Biol 24, 279–279

21. Jang, C. W., Shibata, Y., Starmer, J., Yee, D., and Magnuson, T. (2015) Histone H3.3 maintains genome integrity during mammalian development. Genes & development 29, 1377–1392

22. Ishiuchi, T., Abe, S., Inoue, K., Yeung, W. K. A., Miki, Y., Ogura, A., and Sasaki, H. (2021) Reprogramming of the histone H3.3 landscape in the early mouse embryo. Nat Struct Mol Biol 28

23. Kong, Q. R., Banaszynski, L. A., Geng, F. Q., Zhang, X. L., Zhang, J. M., Zhang, H., O’Neill, C. L., Yan, P. D., Liu, Z. H., Shido, K., Palermo, G. D., Allis, C. D., Rafii, S., Rosenwaks, Z., and Wen, D. C. (2018) Histone variant H3.3-mediated chromatin remodeling is essential for paternal genome activation in mouse preimplantation embryos. Journal of Biological Chemistry 293, 3829–3838

24. Lin, C. J., Conti, M., and Ramalho-Santos, M. (2013) Histone variant H3.3 maintains a decondensed chromatin state essential for mouse preimplantation development. Development 140, 3624–3634

25. Goldberg, A. D., Banaszynski, L. A., Noh, K. M., Lewis, P. W., Elsaesser, S. J., Stadler, S., Dewell, S., Law, M., Guo, X. Y., Li, X., Wen, D. C., Chapgier, A., DeKelver, R. C., Miller, J. C., Lee, Y. L., Boydston, E. A., Holmes, M. C., Gregory, P. D., Greally, J. M., Rafii, S., Yang, C. W., Scambler, P. J., Garrick, D., Gibbons, R. J., Higgs, D. R., Cristea, I. M., Urnov, F. D., Zheng, D. Y., and Allis, C. D. (2010) Distinct Factors Control Histone Variant H3.3 Localization at Specific Genomic Regions. Cell 140, 678–691

26. Wen, D., Noh, K. M., Goldberg, A. D., Allis, C. D., Rosenwaks, Z., Rafii, S., and Banaszynski, L. A. (2014) Genome editing a mouse locus encoding a variant histone, H3.3B, to report on its expression in live animals. Genesis 52, 959–966

27. Banaszynski, L. A., Wen, D., Dewell, S., Whitcomb, S. J., Lin, M., Diaz, N., Elsasser, S. J., Chapgier, A., Goldberg, A. D., Canaani, E., Rafii, S., Zheng, D., and Allis, C. D. (2013) Hira-dependent histone H3.3 deposition facilitates PRC2 recruitment at developmental loci in ES cells. Cell 155, 107–120

28. Liu, X., Wang, C., Liu, W., Li, J., Li, C., Kou, X., Chen, J., Zhao, Y., Gao, H., Wang, H., Zhang, Y., Gao, Y., and Gao, S. (2016) Distinct features of H3K4me3 and H3K27me3 chromatin domains in pre-implantation embryos. Nature 537, 558–562

29. Jones, E. L., Zalensky, A. O., and Zalenskaya, I. A. (2011) Protamine Withdrawal from Human Sperm Nuclei Following Heterologous ICSI into Hamster Oocytes. Protein Peptide Lett 18, 811–816

30. Bui, H. T., Wakayama, S., Mizutani, E., Park, K. K., Kim, J. H., Van Thuan, N., and Wakayama, T. (2011) Essential role of paternal chromatin in the regulation of transcriptional activity during mouse preimplantation development. Reproduction 141, 67–77

31. Bird, A. (2007) Perceptions of epigenetics. Nature 447, 396–398

32. Spivakov, M., and Fisher, A. G. (2007) Epigenetic signatures of stem-cell identity. Nat Rev Genet 8, 263–271

33. Mikkelsen, T. S., Ku, M. C., Jaffe, D. B., Issac, B., Lieberman, E., Giannoukos, G., Alvarez, P., Brockman, W., Kim, T. K., Koche, R. P., Lee, W., Mendenhall, E., O’Donovan, A., Presser, A., Russ, C., Xie, X. H., Meissner, A., Wernig, M., Jaenisch, R., Nusbaum, C., Lander, E. S., and Bernstein, B. E. (2007) Genome-wide maps of chromatin state in pluripotent and lineage-committed cells. Nature 448, 553–U552

34. Zaret, K. S., and Mango, S. E. (2016) Pioneer transcription factors, chromatin dynamics, and cell fate control. Current Opinion in Genetics & Development 37, 76–81

35. Strahl, B. D., and Allis, C. D. (2000) The language of covalent histone modifications. Nature 403, 41–45

36. Jenuwein, T., and Allis, C. D. (2001) Translating the histone code. Science 293, 1074–1080

37. Allis, D. (2002) Translating the histone code: A tale of tails. Molecular biology of the cell 13, 278a–278a

38. Martire, S., Gogate, A. A., Whitmill, A., Tafessu, A., Nguyen, J., Teng, Y. C., Tastemel, M., and Banaszynski, L. A. (2019) Phosphorylation of histone H3.3 at serine 31 promotes p300 activity and enhancer acetylation. Nat Genet 51, 941-+

39. Sato, Y., Hilbert, L., Oda, H., Wan, Y. A., Heddleston, J. M., Chew, T. L., Zaburdaev, V., Keller, P., Lionnet, T., Vastenhouw, N., and Kimura, H. (2019) Histone H3K27 acetylation precedes active transcription during zebrafish zygotic genome activation as revealed by live-cell analysis. Development 146

40. Wang, M., Chen, Z. Y., and Zhang, Y. (2022) CBP/p300 and HDAC activities regulate H3K27 acetylation dynamics and zygotic genome activation in mouse preimplantation embryos. Embo Journal 41

41. Wu, K. L., Fan, D. D., Zhao, H., Liu, Z. B., Hou, Z. Z., Tao, W. R., Yu, G. L., Yuan, S. L., Zhu, X. X., Kang, M. Y., Tian, Y., Chen, Z. J., Liu, J., and Gao, L. (2023) Dynamics of histone acetylation during human early embryogenesis. Cell Discov 9

42. Aoki, F., Worrad, D. M., and Schultz, R. M. (1997) Regulation of transcriptional activity during the first and second cell cycles in the preimplantation mouse embryo. Developmental biology 181, 296–307

43. Yuan, S. L., Zhan, J. H., Zhang, J. Y., Liu, Z. B., Hou, Z. Z., Zhang, C. X., Yi, L. Z., Gao, L., Zhao, H., Chen, Z. J., Liu, J., and Wu, K. L. (2023) Human zygotic genome activation is initiated from paternal genome. Cell Discov 9

44. Chen, Z. Y., and Zhang, Y. (2019) Loss of DUX causes minor defects in zygotic genome activation and is compatible with mouse development. Nat Genet 51, 947-+

45. Armache, A., Yang, S., de Paz, A. M., Robbins, L. E., Durmaz, C., Cheong, J. Q., Ravishankar, A., Daman, A. W., Ahimovic, D. J., Klevorn, T., Yue, Y., Arslan, T., Lin, S., Panchenko, T., Hrit, J., Wang, M., Thudium, S., Garcia, B. A., Korb, E., Armache, K. J., Rothbart, S. B., Hake, S. B., Allis, C. D., Li, H. T., and Josefowicz, S. Z. (2020) Histone H3.3 phosphorylation amplifies stimulation-induced transcription. Nature 583, 852-+

46. Chang, F. T. M., Chan, F. L., McGhie, J. D. R., Udugama, M., Mayne, L., Collas, P., Mann, J. R., and Wong, L. H. (2015) CHK1-driven histone H3.3 serine 31 phosphorylation is important for chromatin maintenance and cell survival in human ALT cancer cells. Nucleic acids research 43, 2603–2614

47. Abe, K., Yamamoto, R., Franke, V., Cao, M. J., Suzuki, Y., Suzuki, M. G., Vlahovicek, K., Svoboda, P., Schultz, R. M., and Aoki, F. (2015) The first murine zygotic transcription is promiscuous and uncoupled from splicing and 3 ’ processing. Embo Journal 34, 1523–1537

48. Majumder, S., Miranda, M., and Depamphilis, M. (1993) Analysis of Gene-Expression in Mouse Preimplantation Embryos Demonstrates That the Primary Role of Enhancers Is to Relieve Repression of Promoters (Vol 12, Pg 1131, 1993). Embo Journal 12, 4042–4042

49. Wiekowski, M., Miranda, M., and Depamphilis, M. L. (1993) Requirements for Promoter Activity in Mouse Oocytes and Embryos Distinguish Paternal Pronuclei from Maternal and Zygotic Nuclei. Developmental biology 159, 366–378

50. Erkek, S., Hisano, M., Liang, C. Y., Gill, M., Murr, R., Dieker, J., Schubeler, D., van der Vlag, J., Stadler, M. B., and Peters, A. H. F. M. (2013) Molecular determinants of nucleosome retention at CpG-rich sequences in mouse spermatozoa (vol 20, pg 868, 2013). Nat Struct Mol Biol 20, 1236–1236

51. Ihara, M., Meyer-Ficca, M. L., Leu, N. A., Rao, S., Li, F., Gregory, B. D., Zalenskaya, I. A., Schultz, R. M., and Meyer, R. G. (2014) Paternal Poly (ADP-ribose) Metabolism Modulates Retention of Inheritable Sperm Histones and Early Embryonic Gene Expression. Plos Genet 10

52. Loppin, B., and Berger, F. (2020) Histone Variants: The Nexus of Developmental Decisions and Epigenetic Memory. Annu Rev Genet 54, 121–149

53. Creyghton, M. P., Cheng, A. W., Welstead, G. G., Kooistra, T., Carey, B. W., Steine, E. J., Hanna, J., Lodato, M. A., Frampton, G. M., Sharp, P. A., Boyer, L. A., Young, R. A., and Jaenisch, R. (2010) Histone H3K27ac separates active from poised enhancers and predicts developmental state. Proceedings of the National Academy of Sciences of the United States of America 107, 21931–21936

54. Li, J., Zhang, J., Hou, W., Yang, X., Liu, X., Zhang, Y., Gao, M., Zong, M., Dong, Z., Liu, Z., Shen, J., Cong, W., Ding, C., Gao, S., Huang, G., and Kong, Q. (2022) Metabolic control of histone acetylation for precise and timely regulation of minor ZGA in early mammalian embryos. Cell Discov 8, 96

55. Wang, M., Chen, Z., and Zhang, Y. (2022) CBP/p300 and HDAC activities regulate H3K27 acetylation dynamics and zygotic genome activation in mouse preimplantation embryos. EMBO J 41, e112012

56. Wu, K., Fan, D., Zhao, H., Liu, Z., Hou, Z., Tao, W., Yu, G., Yuan, S., Zhu, X., Kang, M., Tian, Y., Chen, Z. J., Liu, J., and Gao, L. (2023) Dynamics of histone acetylation during human early embryogenesis. Cell Discov 9, 29

57. Xu, Q., Xiang, Y., Wang, Q., Wang, L., Brind’Amour, J., Bogutz, A. B., Zhang, Y., Zhang, B., Yu, G., Xia, W., Du, Z., Huang, C., Ma, J., Zheng, H., Li, Y., Liu, C., Walker, C. L., Jonasch, E., Lefebvre, L., Wu, M., Lorincz, M. C., Li, W., Li, L., and Xie, W. (2019) SETD2 regulates the maternal epigenome, genomic imprinting and embryonic development. Nat Genet 51, 844–856

58. Wang, C., Liu, X., Gao, Y., Yang, L., Li, C., Liu, W., Chen, C., Kou, X., Zhao, Y., Chen, J., Wang, Y., Le, R., Wang, H., Duan, T., Zhang, Y., and Gao, S. (2018) Reprogramming of H3K9me3-dependent heterochromatin during mammalian embryo development. Nat Cell Biol 20, 620–631

59. Jung, Y. H., Sauria, M. E. G., Lyu, X., Cheema, M. S., Ausio, J., Taylor, J., and Corces, V. G. (2017) Chromatin States in Mouse Sperm Correlate with Embryonic and Adult Regulatory Landscapes. Cell Rep 18, 1366–1382

